# Germline genetic contribution to the immune landscape of cancer

**DOI:** 10.1101/2020.01.30.926527

**Authors:** Rosalyn W. Sayaman, Mohamad Saad, Vésteinn Thorsson, Wouter Hendrickx, Jessica Roelands, Younes Mokrab, Farshad Farshidfar, Tomas Kirchhoff, Randy F. Sweis, Oliver F. Bathe, Eduard Porta-Pardo, Michael J. Campbell, Cynthia Stretch, Donglei Hu, Scott Huntsman, Rebecca E. Graff, Najeeb Syed, Laszlo Radvanyi, Simon Shelley, Denise Wolf, Francesco M. Marincola, Michele Ceccarelli, Jérôme Galon, Elad Ziv, Davide Bedognetti

**Affiliations:** Department of Population Sciences, Beckman Research Institute, City of Hope Comprehensive Cancer Center, Duarte, CA 91010, USA; Department of Laboratory Medicine, Helen Diller Family Comprehensive Cancer Center, University of California, San Francisco, San Francisco, CA 94143, USA; Qatar Computing Research Institute, Hamad Bin Khalifa University, Doha, Qatar; Institute for Systems Biology, Seattle, WA 98109, USA; Research Branch, Sidra Medicine, PO Box 26999 Doha, Qatar; Department of Surgery, Leiden University Medical Center, 2333 ZA Leiden, the Netherlands; Department of Oncology, University of Calgary, Alberta AB T2N 4N1, Canada; Arnie Charbonneau Cancer Institute, Calgary, Alberta AB T2N 4N1, Canada; Department of Biomedical Data Science and Institute for Stem Cell Biology and Regenerative Medicine, School of Medicine, Stanford University, Stanford, CA 94305, USA; Perlmutter Comprehensive Cancer Center, New York University School of Medicine, New York University Langone Health New York, New York, NY 10016, USA; Department of Medicine, Section of Hematology/Oncology, Committee on Clinical Pharmacology and Pharmacogenomics, Committee on Immunology, University of Chicago, Chicago, IL 60637, USA; Department of Surgery, University of Calgary, Calgary, Alberta AB T2N 4N1, Canada; Barcelona Supercomputing Center (BSC), Josep Carreras Leukaemia Research Institute (IJC), Badalona, 08034 Barcelona, Catalonia, Spain; Department of Surgery, University of California, San Francisco, San Francisco, CA 94143, USA; Department of Medicine, Institute for Human Genetics, Helen Diller Family Comprehensive Cancer Center, University of California, San Francisco, San Francisco, CA 94143, USA; Department of Epidemiology and Biostatistics, University of California, San Francisco, San Francisco, CA 94143, USA; Ontario Institute for Cancer Research, Toronto, Ontario M5G 0A3, Canada; Department of Research and Development, Leukemia Therapeutics, LLC, Hull, MA 02045, USA; Refuge Biotechnologies Inc., Menlo Park, CA 94025, USA; Department of Electrical Engineering and Information Technology, University of Naples “Federico II”, 80128 Naples, Italy; Istituto di Ricerche Genetiche “G. Salvatore”, Biogem s.c.ar.l, 83031 Ariano Irpino, Italy; INSERM, Laboratory of Integrative Cancer Immunology, Sorbonne Université, Sorbonne Paris Cité, Université Paris Descartes, Université Paris Diderot, Centre de Recherche des Cordeliers, 75006 Paris, France; College of Health and Life Sciences, Hamad Bin Khalifa University, Doha, Qatar

## Abstract

The role of germline genetics in shaping the tumor immune landscape is largely unknown. Using genotypes from >9,000 individuals in The Cancer Genome Atlas, we investigated the association of common and rare variants with 139 well-defined immune traits. Our analysis of common variants identified 10 immune traits with significant heritability estimates, and an additional 23 with suggestive heritability, including estimates of T-cell subset abundance and interferon signaling. We performed genome-wide association on the 33 heritable traits and identified 23 genome-wide significant loci associated with at least one immune trait, including SNPs in the *IFIH1* locus previously associated with several autoimmune diseases. We also found significant associations between immune traits and pathogenic or likely-pathogenic rare variants in *BRCA1* and in genes functionally linked to telomere stabilization, and Wnt/Beta-catenin signaling. We conclude that germline genetic variants significantly impact the composition and functional orientation of the tumor immune microenvironment.

## Introduction

Tumor-host interactions are important determinants of carcinogenesis and metastatic progression (Hanahan and Weinberg, 2011). The type and density of immune-cell infiltration in the tumor microenvironment are associated with differential prognosis (Bedognetti et al., 2019; Galon et al., 2006; Pagès et al., 2018). Metastases evolve in the context of selection by the immune system (Angelova et al., 2018). Immunotherapy approaches, including the use of monoclonal antibodies that target immune inhibitory signaling (immune checkpoints), have emerged as the standard of care for many cancers (Wilky, 2019). However, checkpoint inhibitors are only effective in a subset of cancer types, and even among cancer types in which they are active, most patients do not respond to them (Chen and Mellman, 2017; Sweis and Luke, 2017). Given these limitations, substantial effort has been directed at identifying factors modulating immunotherapy responsiveness. Tumor immune infiltration has been shown to predict responsiveness to immunotherapy (Tumeh et al., 2014). Therefore, advancing our understanding of the factors that underlie the host immune response to the tumor may help identify additional pathways to target for therapy (Bedognetti et al., 2019).

Tumor-intrinsic factors are associated with immune infiltration and response to immunotherapy. For example, a higher burden of somatic mutations and a T-cell inflamed phenotype are independently associated with response to immune checkpoint inhibitors (Cristescu et al., 2018; Samstein et al., 2019; Snyder et al., 2014; Trujillo et al., 2018; Van Allen et al., 2015). DNA mismatch repair deficiency is mechanistically linked to intratumoral adaptive immunity, and microsatellite instability (MSI) is associated with response to immunotherapy (Mlecnik et al., 2016). Aneuploidy, which might alter tumor signals required for immune-cell chemoattraction and optimal cytotoxic response, is associated with decreased immune infiltration and a lower likelihood of responding to immunotherapy (Davoli et al., 2017). Specific somatic driver mutations have been correlated with differences in the quantity and types of immune cells infiltrating the tumor and with differences in response to immunotherapy (Riaz et al., 2016; Thorsson et al., 2018). Environmental factors such as the microbiome may also modify the immune microenvironment and affect response to immunotherapy (Fessler et al., 2019). Host genetic factors have not been explored in depth as predictors of immune responsiveness (Bedognetti et al., 2019).

Germline variants exert strong effects on circulating leukocyte counts (Keller et al., 2014) and on leukocyte fractions (Orru et al., 2013). Immune-mediated tissue rejection in organ or bone marrow transplantation (Yang and Sarwal, 2017) and autoimmunity (Ye et al., 2018) have been associated with common germline variations in genome-wide association studies (GWAS). Candidate gene studies of patients treated with cytokine-based therapies or anti-CTLA-4 monoclonal antibodies have linked treatment outcome to polymorphisms of immune-related genes including *CTLA4* (Queirolo et al., 2017)*, IRF5* (Uccellini et al., 2012), and *CCR5* (Bedognetti et al., 2013; Ugurel et al., 2008). Polymorphisms at the *HLA* and Fc loci have also been associated with outcomes in patients treated with checkpoint inhibition (Arce Vargas et al., 2018; Chowell et al., 2018). Furthermore, somatic or germline defects in mismatch-repair genes significantly increase the likelihood of response to checkpoint inhibitors, (Le et al., 2015, 2017a) likely by increasing the neoantigen load of cancer cells (Mandal et al., 2019).

The Cancer Genome Atlas (TCGA) project has dramatically increased our understanding of cancer pathogenesis and allowed the characterization of phenotypic features associated with disease aggressiveness and outcomes (Hutter and Zenklusen, 2018). A recent comprehensive analysis of immune signatures in data from the TCGA identified several important features of immune response that predict survival across many tumor types (Thorsson et al., 2018). Here, in the same cohort, we used a pan-cancer approach to evaluate the contribution of germline variation to anti-tumor immune response. First, we used genotype data from common variants to calculate genome-wide heritability of immune traits. Next, we performed GWAS for immune traits to identify the loci with the strongest effects. Finally, we examined the contribution of rare germline variants, looking at known cancer susceptibility genes and pathways.

## Results

### Overview of the discovery approach

To examine the contribution of germline genetic variation to the functional orientation of the immune microenvironment, we conducted heritability analysis, GWAS (N=9,603), and rare variant analysis (N=9,138) across 30 non-hematological cancer types characterized by the TCGA (**Figure 1 top**). All analyses adjusted for cancer type, age at diagnosis, sex, and the first seven components from principal component analysis (PCA) done on SNP data, which largely capture genetic ancestry (see Methods).

**Figure 1.**
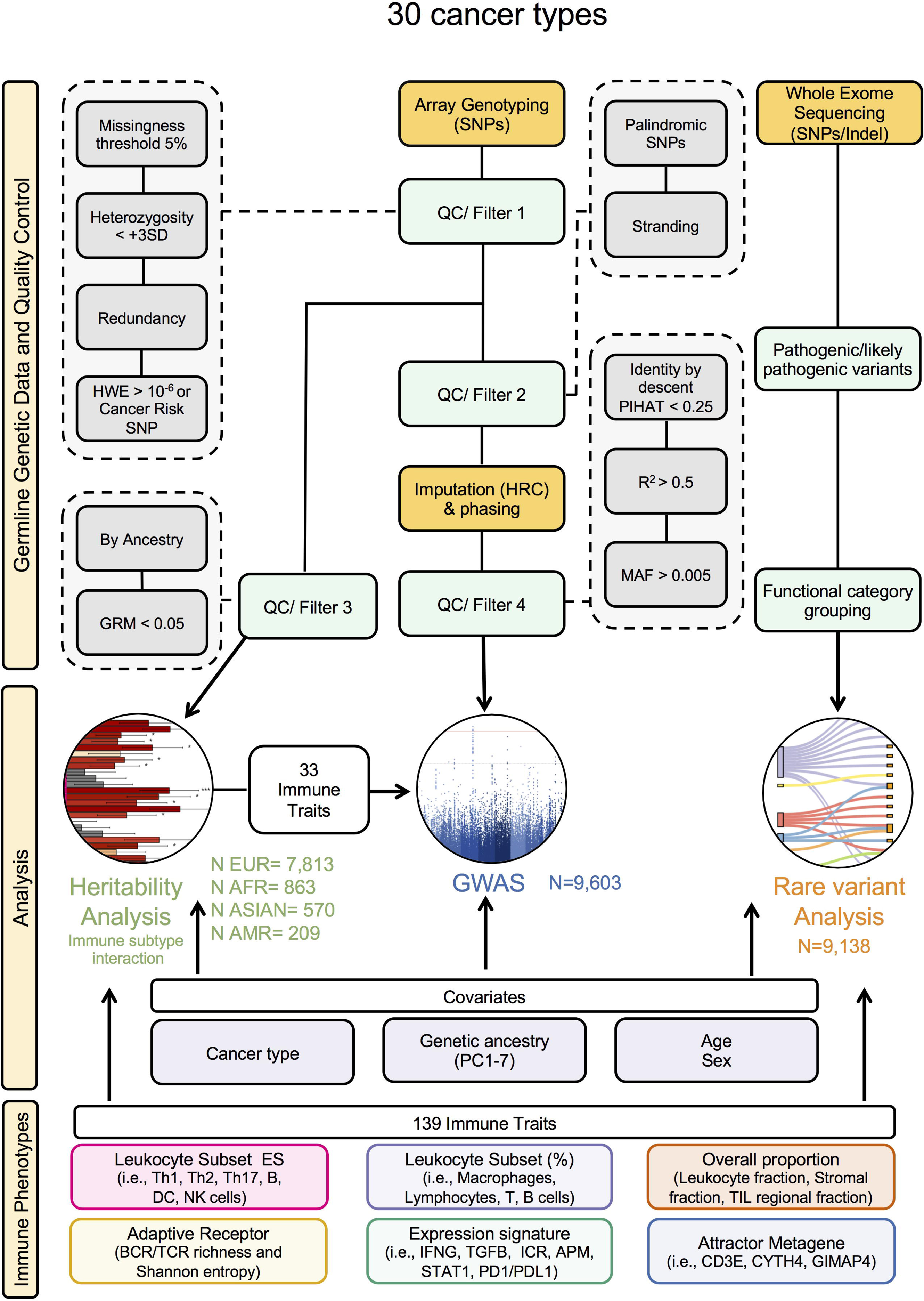
Overview of the discovery approach. Flowchart showing analytic workflows, source of genetic germline data, quality control filtering, and immune traits used in the analysis.

We considered 139 well-characterized immune traits estimated in the previously described TCGA Immune analysis (Thorsson et al., 2018) (**Figure 1 bottom panel**). We divided the traits into 6 categories based on the approach used to derive them and what they were intended to detect: (1) leukocytes subset enrichment score (LSES) and (2) leukocyte Subset proportion (%), estimating the leukocyte subpopulations abundancy within the tumor and their proportion within infiltrating leukocytes, respectively, (3) Overall proportion, which include measures of leukocyte infiltration and stromal contents, (4) Adaptive receptor, quantifying TCR and BCR diversity (TCR and BCR Shannon entropy and richness) (5) expression signatures, consisting in a collection of annotated functional signatures summarizing different immune-related biological process (e.g., wound healing, Interferon (IFN) and TGF-β signaling, antigen-presenting machinery etc.), and (6) Attractor Metagene, which includes co-expression signatures (metagene attractors) derived from the whole TCGA dataset. All of the immune traits were derived from RNA-seq, with the exception of overall proportion estimates (derived from H&E tissue imaging and methylation arrays).

The 139 traits were clustered based on their Pearson correlation coefficients, and 6 groups of highly correlated traits were defined and referred to here as “modules” (**Fig S1**). The largest group included expression signatures that were highly correlated with leukocyte fraction and lymphocyte infiltration estimates (Lymphocyte infiltration module). A group of traits capturing monocyte-function and macrophages infiltration formed a second module that was highly correlated with the lymphocyte infiltration module (Monocyte/macrophage module). This module also included several MHC-related expression signatures. Traits capturing interferon signaling were highly correlated and formed a third distinct module (IFN module). The next two modules included traits associated with TGF-β signaling (TGF-β module) and wound healing (wound healing module), respectively. The last module mostly included leukocyte subsets such as T helper cells subset, CD8 cytotoxic and natural killer (NK) cells (Cytotoxic module). Although most traits fit within these modules, a subset (N=42), did not cluster within any module. In addition, there was substantial correlation between traits across modules, particularly between the lymphocyte infiltration and monocyte/macrophage modules and between both of these and the IFN modules. A subset of traits clustering within the Cytotoxic module were significantly correlated with the IFN, lymphocyte infiltration, and monocyte/macrophage modules.

GWAS were only performed on traits that demonstrated significant heritability, since these were most likely to have significant genetic effects. For rare variant analyses, we examined all 139 traits, and focused on well-annotated pathogenic or likely mutations (Huang et al., 2018) within high penetrance cancer susceptibility genes.

### Genome-wide heritability of immune traits

Genome-wide heritability is the fraction of phenotypic variance explained by genetic variants (Zaitlen and Kraft, 2012). It can be estimated using common variants based on the relationship between genome-wide genotypic correlation and phenotypic correlation among unrelated individuals (Yang et al., 2010, 2011). We performed heritability analysis on the 139 traits using a mixed-model approach implemented in GREML (see Methods). Heritability analyses were conducted separately within each ancestral subgroup (N_European_=7,813, N_African_=863, N_Asian_=570, and N_American_=209 individuals), which were derived from ancestry analysis using the genotype data (**Figure S2A-B**).

In the pan-cancer analysis, we found 10 immune traits with significant heritability after correction for multiple comparison testing (FDR *p* < 0.05), and 23 other traits with nominally significant heritability (*p* < 0.05) in at least one ancestry group (**Figure 2A, Figure S2C**). Within the European ancestry group, 28 traits were at least nominally significantly heritable. The most heritable traits, with ∼15-20% heritability and FDR *p* < 0.05, were members of the Cytotoxic module (**Figure S1A**) and represent T cell subsets estimated using LSES including CD8 T cells, T helper cells, T follicular helper cells (Tfh), T effector memory (Tem) cells, T central memory (Tcm) cells, NK cells and eosinophils (**Figure 2A**). An Antigen Presenting Machinery signature (APM1) clustered within the Cytotoxic module (**Figure S1A**) and was also highly heritable (∼20% heritability and FDR *p* < 0.05). T helper 1 (Th1) cells LSES, which were also part of this module (**Figure S1**), showed lower but nominally significant heritability (**Figure 2A**). The second most heritable traits, with ∼15% heritability, included highly correlated IFN-related signatures (Interferon Cluster 21214954, GP11 Immune IFN, IFN 21978456, Module3 IFN score, IFIT3) and activated dendritic cells (aDC), which cluster within the IFN module (**Figure S1A**). Among these, Interferon Cluster 21214954 and aDC LSES were significant after correction for multiple hypothesis testing. In addition, we detected nominally significant heritability for Th2 cells LSES and Th17 cells LSES, proportion of T cells CD8+, memory B cells, and eosinophils within leukocytes (Leukocyte Subset %). The CD8+ T-cell/CD68+ ratio, a B-cell metagene score (B-cell mg IGJ), macrophages LSES, neutrophils LSES, and other signatures belonging to the macrophage/monocyte module, including major histocompatibility complex class-II (MHC2 21978456) and Siglec-regulation (G SIGLEC9), an attractor metagene, showed nominally significant heritability.

**Figure 2.**
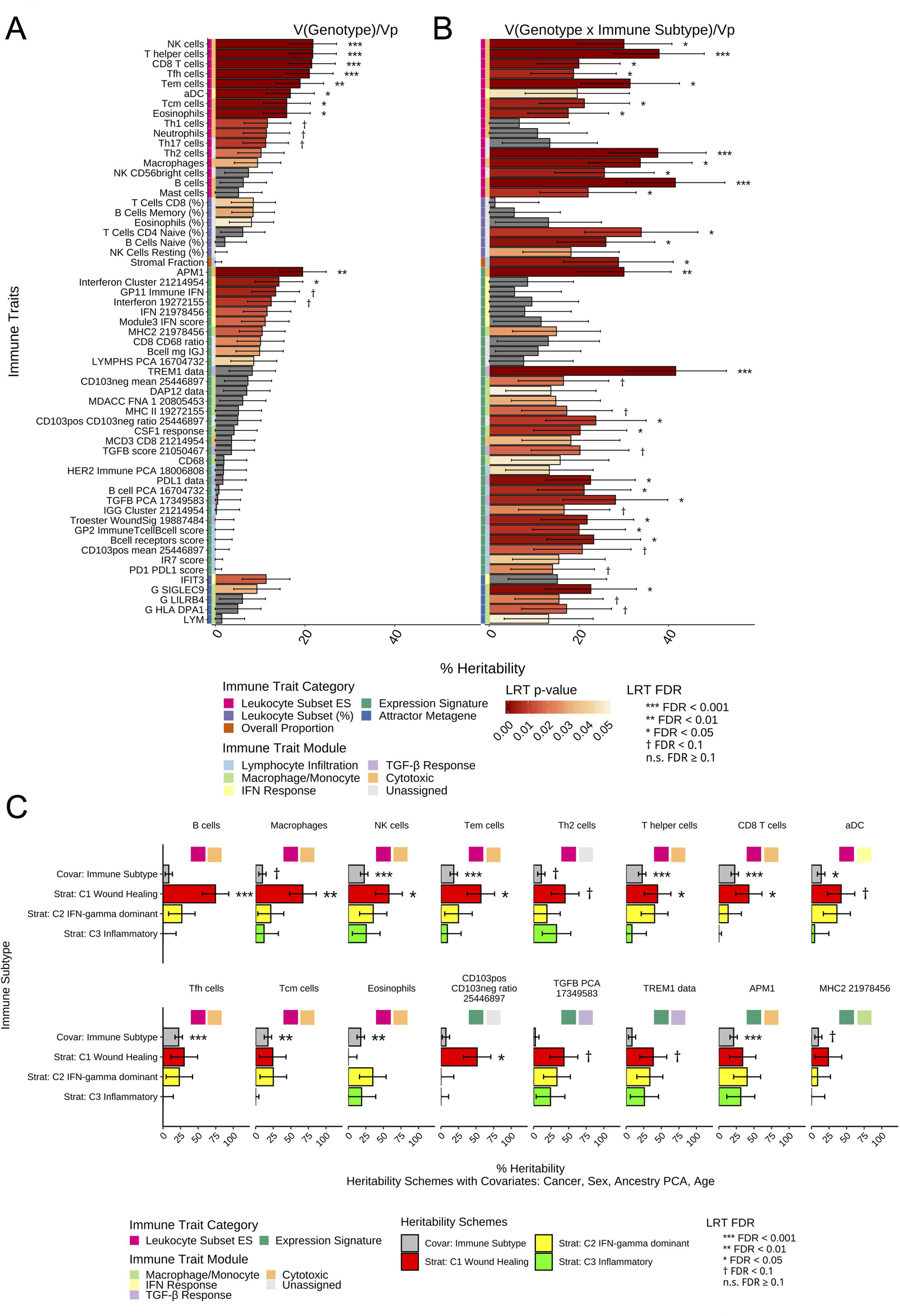
Genome-wide heritability of immune traits. Genome-wide complex trait analysis (GCTA) using genomic-relatedness-based restricted maximum-likelihood (GREML) considers the simultaneous effect of all SNPs and provides estimates of the proportion of phenotypic variance explained by the genetic variance, V(Genotype)/Vp. (**A**) Percentage of variance explained by common genetic variants. Twenty-eight of 139 immune traits analyzed in the European ancestry group (n=7,813) show nominally significant level of genome-wide heritability (LRT *p* < 0.05), with 10 traits (FDR < 0.05) and 15 traits (FDR < 0.1) showing significant heritability after correction for multiple hypothesis testing. These immune traits include measures of leukocyte subset enrichment score (ES) and percentages (%), as well as immunomodulatory expression signatures and attractor metagenes. (**B**) Percentage of variance of immune traits accounted for by interaction between germline genotypes and immune subtypes (G x Immune Subtype). In the subset of individuals with immune subtype information (n = 6,586), 44 immune traits show nominally significant heritability of interaction effects, V(GxImmune Subtype)/Vp (LRT *p* < 0.05), and 26 traits show significant heritability of interaction effects (FDR < 0.05). Corresponding heritability estimates with standard errors are shown in (**A**) and (**B**) for each of the 59 immune traits identified with immune trait categories and corresponding immune trait modules annotated by the colorbar. (**C**) Immune subtype-specific heritability analysis conducted for immune traits with significant G x Immune Subtype interaction demonstrate the differential effect of immune subtypes on the contributions of genotypic variance to overall phenotypic variance. Heritability was calculated in three of the six immune subtype groups with sufficient cohort size: C1 Wound Healing (n=1752), C2 IFN-γ dominant (n=1813), and C3 Inflammatory (n=1737), as well as with immune subtype as an additional covariate. Stratified analysis of the 44 traits with at least nominally significant G x Immune Subtype interaction effects show 16 traits with significant V(G)/Vp heritability in at least one of the immune subtypes or with immune subtype as a covariate (LRT FDR < 0.1). Heritability analysis are are run with age, tumor type, sex and PC1-7 as covariates.

Despite the limited cohort size, specific immune traits were also found to be heritable within the African and Asian ancestry groups, though they were only nominally significant. Within the African ancestry group, NK CD56dim LSES and Cytotoxic cell LSES, single gene immune therapy target *PDCD1* expression (PD1 data), and TGF-β immunomodulatory signaling (TGFB PCA 17349583), showed nominally significant heritability. For individuals of Asian ancestry, Tcm LSES and the proportion of NK cells activated (%) showed nominally significant heritability (**Figure S2C**).

### Variation of Heritability of Immune Traits across Immune Subtypes

Since there was considerable heterogeneity between tumor types, we investigated whether heritability varies among tumor immune subtypes defined by a computational analysis of the expression pattern of immunological parameters (Thorsson et al., 2018). These six distinct immune subtypes include: wound healing (C1), IFN-γ dominant (C2), inflammatory (C3), lymphocyte depleted (C4), immunologically quiet (C5), and TGF-β dominant (C6) (**Figure S1B**). These immune subtypes formed larger subsets than any of the individual tumor types, facilitating heritability analyses that generally require large numbers (Visscher et al., 2014). We performed statistical interaction analysis using GREML, which determined the fraction of the phenotypic variance, Vp, contributed by the interaction between genotype and immune subtype, V(Genotype x Immune Subtype)/Vp (**Figure 2B**) in the European ancestry group. We found 26 immune traits with significant variance of genotype-immune subtype interaction effects at FDR *p* < 0.05, and 18 other nominally significant traits (*p* < 0.05). These interactions suggest that the contribution of genotype to phenotypic variation of immune traits differs among immune subtypes. We found substantial overlap between immune traits with nominally significant heritability and with interaction effects. Eight of the heritable traits from the Cytotoxic Module had significant interactions with immune subtype including: LSES of CD8 T cells, T helper cells, T Tfh cells, Tem cells, Tcm cells, NK cells, eosinophils and APM1 expression signature. In addition, LSES of NK CD56 bright cells, B cells, and mast cells show significant interaction effects of genotype-immune subtype. The majority of immune traits with significant and nominally significant genotype-immune subtype interaction were those associated with the macrophage/monocyte, overall lymphocyte infiltration, and TGF-β response module. On the other hand, the six immune traits associated with the IFN module showed no genotype-immune subtype interaction, suggesting that the genotypic contribution to phenotypic variation of these traits was consistent across immune subtypes.

To understand how immune subtype influences heritability of the 44 immune traits with at least nominally significant genotype-immune subtype interaction effects (*p* < 0.05), we performed heritability analyses stratified by immune subtype. We performed these only in subtypes with sufficient sample size: C1 Wound healing (N_C1_=1,752), C2 IFN-γ dominant (N_C2_=1813), C3 Inflammatory (N_C3_=1737). C4 Lymphocyte depleted, C5 Immunologically quiet and C6 TGF-β dominant immune subtypes were insufficiently powered for this analysis. We found significant heritability estimates (FDR from *p* < 0.1 to *p* < 0.001 significance levels) largely within the C1 Wound healing immune subtype, but not within the more immune-active subtypes C2 IFN-γ dominant and C3 Inflammatory (**Figure 2C**). Highly heritable (FDR *p* < 0.1) immune traits within C1 Wound healing included: LSES of B Cells, macrophages, NK cells, Tem cells, T helper cells, CD8 T cells (Cytotoxic module), as well as aDC (IFN module) and Th2 cells; expression signatures TREM1 and TGFB PCA (TGF-β response module); and CD103+/CD103-ratio (**Figure 2C**).

### Genome-wide association for variants affecting immune traits

We selected the 33 immune traits with nominally significant heritability (*p* < 0.05) in at least one ancestry group to perform GWAS. We identified 598 genome-wide significant (*p* < 5×10^−8^) associations at 23 loci for 10 immune traits. We also identified an additional 1,196 suggestive (*p* < 1×10^−6^) associations for 33 traits (**Figure 3A**). These loci are annotated by genes within +/-50kb of the SNP or by rsID when no gene is in proximity in the combined Manhattan plot (**Figure 3A**).

**Figure 3.**
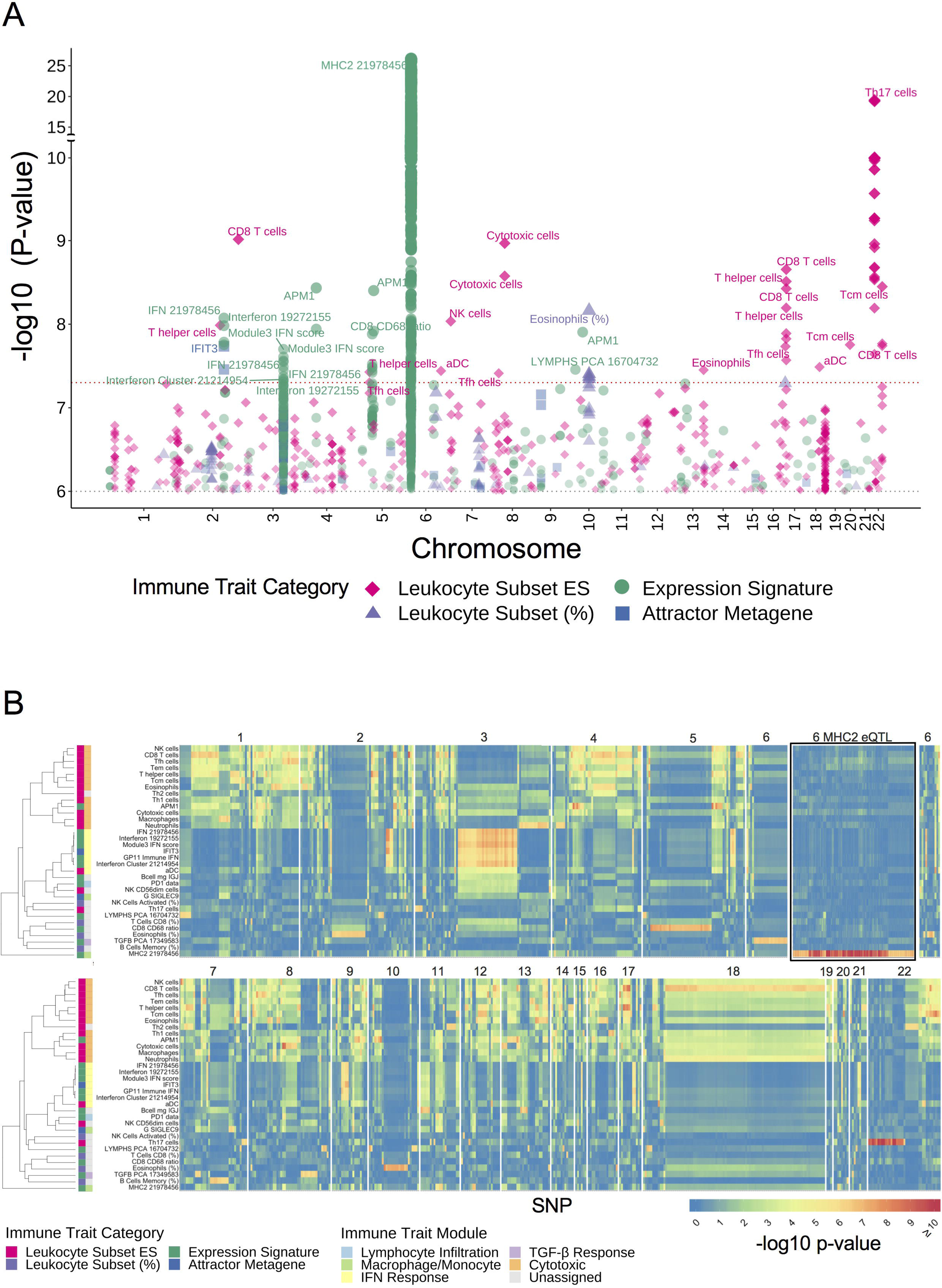
Genome-wide associations for variants affecting immune traits. Genome-wide association studies (GWAS) performed on the 33 immune traits showing genome-wide heritability in the ancestry clusters identifies 23 loci with 598 genome-wide significant associations between single SNPs and immune disposition in 10 immune traits (*p* < 5×10^−8^), and an additional 1,196 suggestive associations in 33 traits (*p* < 1×10^−6^). (**A**) Combined Manhattan plot representing -log10 *p* of the significant and suggestive GWAS hits by chromosomal position across the 33 immune traits encompassing four phenotypic categories: leukocyte subset enrichment score (ES) and percentage (%), immunomodulatory expression signature and attractor metagene. Of the 598 genome-wide significant associations, 584 are represented by unique SNPs, 10 of which have multiple hits in at least 2 traits; of the 1,196 suggestive associations, 1,022 are represented by unique SNPs, 70 of which have multiple hits in at least 2 traits. (**B**) Heatmap demonstrating pleiotropy of the top associations across 33 immune traits. The heatmap is facetted by chr showing the GWAS -log10 *p* of the 1,587 significant and suggestive SNPs across each of the 33 immune subtypes. Immune traits are clustered based on the Pearson correlation of the GWAS -log10 *p* with immune trait categories and corresponding immune trait modules annotated by the colorbar Two eQTL regions are identified, one on chr 6 for the MHC2 signature with 864 SNPs at the *HLA* loci (shown condensed), and one on chr 22 for Th-17 cells with 32 SNPs at the *IL17R* locus. All GWAS are are run with age, tumor type, sex and PC1-7 as covariates.

Two of the 23 loci with the strongest associations (*p* < 1×10^−10^ to *p* < 1×10^−25^) included SNPs that map to +/-50 KB (or 1 MB in the case of HLA) from genes that are part of the signature of the associated immune trait. These included SNPs at the *HLA* locus, which are associated with the MHC2 expression signature, and SNPs at the *IL17R* locus, which make up the Th17 cell enrichment score. We concluded that these SNP associations represented simple expression quantitative trait loci (eQTLs) and we did not consider them further.

In contrast, the remaining 21 loci were not near the genes comprising the associated signatures. Thus, they likely represent SNPs affecting the overall immune trait. The majority of these 21 loci are associated with leukocyte subset enrichment and interferon signaling. Of the 598 genome-wide significant associations, 584 were represented by unique SNPs, 10 of which had significant association with at least 2 traits. Among the 1,196 suggestive associations, 1,022 were represented by unique SNPs, 70 of which had multiple suggestive hits in at least two traits, suggesting pleiotropic associations of significant SNPs.

To examine pleiotropy, we clustered all 33 immune traits based on the association *p* values of all significant and suggestive SNPs found to be associated with at least one trait (**Figure S3B**). The results generally recapitulated clustering based on the correlation of their phenotypic values (**Figure S3A-B**). To better understand the overall genomic context of SNPs associated with immune-associated traits in cancer, we visualized the associations with a heatmap (**Figure 3B**). Traits with shared associations (adjacent rows) tended to have similar overall expression. SNPs that were associated with one of the T-cell LSES tended to be associated with multiple other T-cell subsets. For example, significant and suggestive SNPs in chromosome (chr) 13 associated with CD8 T-cells that were part of the Cytotoxic module showed weaker associations (*p* < 1×10^−4^ to *p* < 1×10^−6^) across a number of other traits from the Cytotoxic module (magenta), and largely no associations across other traits; this phenomenon was consistent across other chromosomes (chrs) for traits within this module. SNPs associated with one of the traits from the IFN module were usually associated with the other traits within that module, but distinct from Cytotoxic module traits. For example, SNPs at chr 2 were associated with the IFN module traits, but not other traits. However, a few of the top loci were associated with traits from both IFN and Cytotoxic modules. For example, significant SNPs in chr 3 associated with IFN module immune traits showed weaker associations with other immune traits, including eosinophils, Th1 cells and cytotoxic T-cells within the Cytotoxic module. Finally, simple eQTLs for MHC2 in chr 6 and Th17 LSES (driven by *IL17R* expression) on chr 22 showed largely no association outside of their respective associated traits.

### Genetic variants associated with differential Interferon signaling disposition and discovery of association with *IFIH1*, an autoimmune-related locus

We found two loci associated with traits from the interferon (IFN) module one on chr 2 and another on chr 3, as shown in the LocusZoom and Manhattan plot for one of the IFN-related traits (**Figure 4A and Figure S4A**). The locus on chr 2 is represented by SNPs rs2111485 and rs1990760, which were both significantly associated with three of the IFN module traits (**Figure 4A and Figure S4A-E**) and had suggestive associations with two other IFN module traits. These two SNPs map to the *IFIH1* gene, and are in high linkage disequilibrium (LD) with each other (r^2^ > 0.8) (**Figure 4B**). Two additional SNPs, rs17716942 and rs13406089, map to the *IFIH1-GCA-KCNH7* locus, but have lower LD r^2^ values (r^2^ < 0.5). Rs2111485, rs1990760, and rs17716942, have been reported to be associated with a number of autoimmune diseases such as psoriasis, vitiligo, systemic lupus erythematosus, ulcerative colitis, inflammatory bowel disease, Crohn’s disease, and type I diabetes mellitus in the GWAS catalog (Buniello et al., 2019) (**Figure 4C**). The direction of the effects of the top SNP was largely consistent across tumor types as seen in the forest plot (**Figure 4D**), while the two SNPs with lower LD had a more variable effect across tumor types.

**Figure 4.**
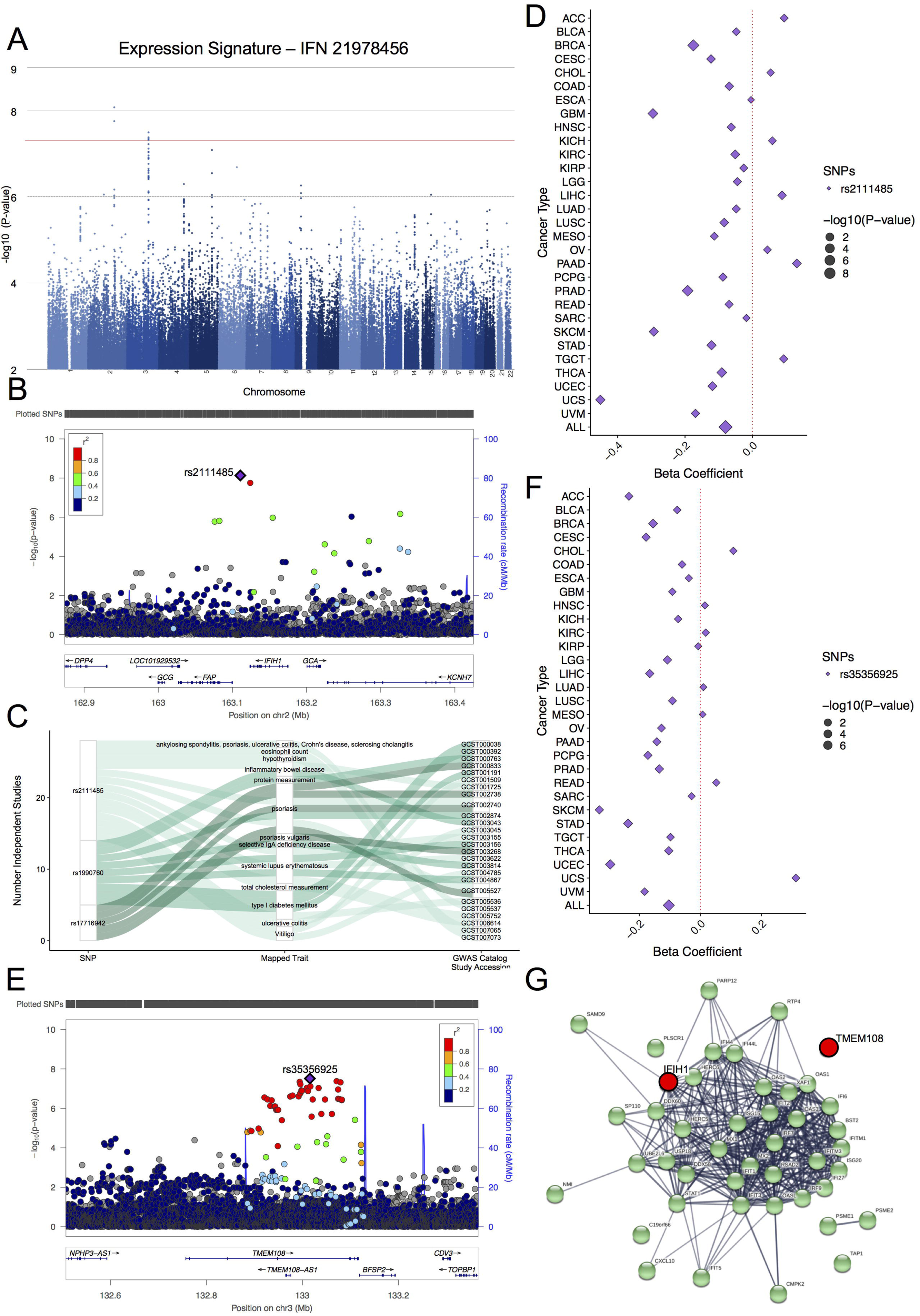
Genetic variants associated with differential Interferon signaling disposition. Genome-wide association studies (GWAS) identifies 17 associations between seven SNPs and five Interferon signatures reaching genome-wide significance (*p* < 5×10^−8^), and additional 152 suggestive associations between 29 SNPs and 6 interferon-related expression signatures (*p <* 1×10^−6^). The pancancer GWAS are run with age, tumor type, sex and PC1-7 as covariates. (**A**) Manhattan plot of GWAS -log10 *p* for a representative Interferon signature, IFN 21978456, shows two main GWAS peaks, one on chr 2 and another on chr 3. (**B**) Locus zoom plot of the association results maps the genomic location of the two significant and two suggestive SNPs on chr 2 to the *IFIH1-GCA-KCNH7* locus. Linkage disequilibrium estimates (r^2^ color map) and recombination rates around the genome-wide significant loci (blue line) are shown. (**C**) Three of the SNPs in the *IFIH1-GCA-KCNH7* locus have been mapped to 13 traits which are predominantly autoimmune-related in 26 independent studies in the GWAS Catalog (Buniello A, et al., *Nucleic Acids Res*, 2019). (**D**) Forest plot of Beta coefficients from association tests within each cancer type compared to the pan-caner result for the most significant GWAS SNP, rs2111485. Within cancer association tests are run with age, sex and PC1-7 as covariates, except in CESC, OV, PRAD, TGCT, UCEC and UCS where only age and PC1-7 are used. (**E**) Locus zoom plot maps the genomic location of the 5 significant and 32 suggestive SNPs on chr 3 to the *TMEM108* locus. (**F**) Forest plot of Beta coefficients of association tests within each cancer type compared to the pan-cancer result for the most significant GWAS SNP, rs35356925. **(G)** Protein-protein interaction network (String-db, minimum interaction score confidence > 0.7) between the 43 genes incorporated into the 6 Interferon-related signatures, and *IFIH1* and *TMEM108*. ACC: Adrenocortical Carcinoma; BLCA: Bladder Urothelial Carcinoma; BRCA: Breast Invasive Carcinoma; CESC: Cervical Squamous Cell Carcinoma and Endocervical Adenocarcinoma; CHOL: Cholangiocarcinoma; COAD: Colon Adenocarcinoma; ESCA: Esophageal Carcinoma; GBM: Glioblastoma; HNSC: Head and Neck Squamous Cell Carcinoma; KICH: Kidney Chromophobe; KIRC: Kidney Renal Clear Cell Carcinoma; KIRP: Kidney Renal Papillary Cell Carcinoma; LGG: Low Grade Glioma; LIHC: Liver Hepatocellular Carcinoma; LUAD: Lung Adenocarcinoma; LUSC: Lung Squamous Cell Carcinoma; MESO: Mesothelioma; OV: Ovarian Serous Cystadenocarcinoma; PAAD: Pancreatic Adenocarcinoma; PCPG: Pheochromocytoma and Paraganglioma; PRAD: Prostate Adenocarcinoma; READ: Rectum Adenocarcinoma; SARC: Sarcoma; SKCM: Skin Cutaneous Melanoma; STAD: Stomach Adenocarcinoma; TGCT: Testicular Germ Cell Tumors; THCA: Thyroid Carcinoma; UCEC: Uterine Corpus Endometrial Carcinoma; UCS: Uterine carcinosarcoma; UVM Uveal Melanoma; ALL: All 30 Cancers combined.

The locus on chr 3 included six genome-wide significant SNPs associated with three of the six traits from the IFN module. These SNPs map to the *TMEM108* gene, which also overlaps with *TMEM108-AS1* Antisense RNA1 gene, and are in high LD with each other (r^2^ > 0.8) (**Figure 4E**). Due to their high LD, the effects of the SNPs were consistent with each other, and we only show the top SNP in the forest plot, where the effect of the SNP within each tumor is largely consistent with the pan-cancer analysis (**Figure 4F**).

Based on known protein-protein interactions (PPI), as annotated in String database (minimum interaction score confidence > 0.7), we found that *IFIH1* formed a highly connected PPI network with the majority of genes that form the traits that belonged to the IFN module (**Figure 4G**). In contrast, *TMEM108* showed no known direct PPI with the IFN genes. However, as noted above, the SNPs at this locus are also nominally associated with some of the T-cell subsets.

### Genetic variants associated with Cytotoxic module

Fourteen of the genome-wide significant loci were associated with eight distinct traits derived from LSES that cluster within the Cytotoxic module (**Figure 5A inset**). We show each of the genome-wide significant and suggestive SNPs mapping to these 14 loci in a combined Manhattan plot (**Figure 5A**).

**Figure 5.**
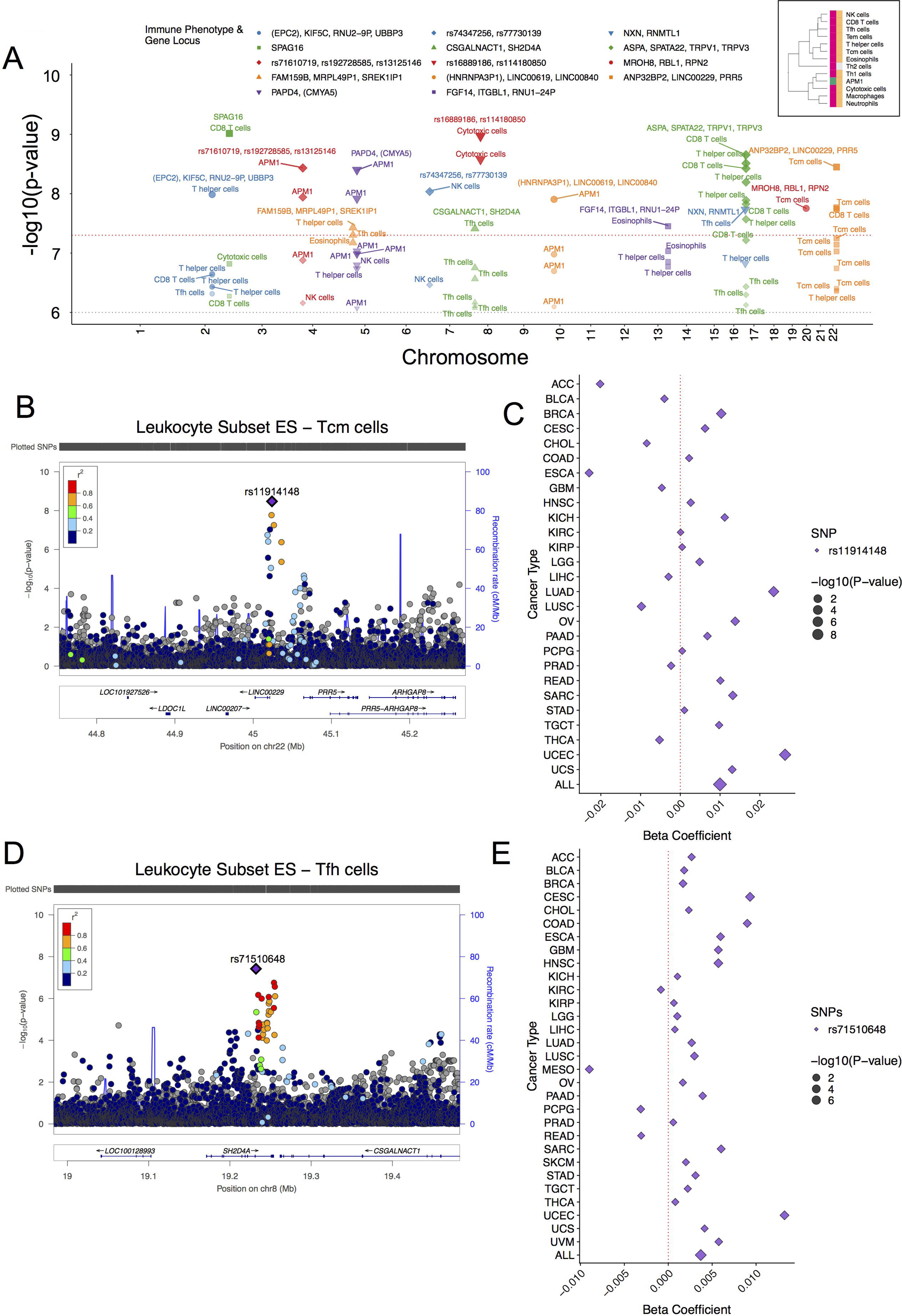
Genetic variants associated with T-cell subset enrichment. Clustering of resulting GWAS -log10 *p* identified highly correlated associations of SNPs across 13 immune traits predominantly associated with T-cell subset enrichment scores: NK, CD8 T, T-follicular helper, T-effector memory, T-helper, T-central memory, T-helper 1, T-helper 2, and cytotoxic cells, along with cells of innate immunity eosinophils, macrophages and neutrophils, and an enrichment score for antigen presenting machinery (APM1) (inset). The pancancer GWAS are run with age, tumor type, sex and PC1-7 as covariates. (**A**) Combined Manhattan plot showing 14 distinct loci and 26 genome-wide significant associations (*p* < 5×10^−8^) between 22 SNPs and eight immune traits within the T-cell subset-dominant cluster: T helper, CD8 T, T-follicular helper, T-central memory, cytotoxic, and NK cells, eosinophils and APM1. Each locus is defined to be +/-50KB region around each SNP. Of these, 13 loci have multiple significant or suggestive (*p* < 1×10^−6^) hits, and 10 have hits in the same region in more than one immune trait. (**B**) Ten associations involving 8 SNPs on chr 22 and T-central memory, CD8 T, and T helper cells mapping to the *LINC00229* locus are represented in a locus zoom plot. (**C**) Forest plot of Beta coefficients from association tests within each cancer type compared to the pan-cancer result for SNP rs11914148 in T-central memory cells. (**D**) Six associations involving 6 SNPs on chr 8 and T-follicular helper cells mapping to the *SH2D4A* locus are represented in a locus zoom plot. (**E**) Forest plot of Beta coefficients from association tests within each cancer type compared to the pan-cancer result for SNP rs71510648 in T-follicular helper cells. Within cancer association tests are run with age, sex and PC1-7 as covariates, except in CESC, OV, PRAD, TGCT, UCEC and UCS where only age and PC1-7 are used.

The locus on chr 22 includes 3 SNPs with genome-wide significant associations including two SNPs, rs73889576 and rs11914148, associated with Tcm cells LSES (**Figure 5B**). A third SNP, rs572393792, with a suggestive association with Tcm cells LSES, had a genome-wide significant association with CD8 T cells LSES and suggestive association with T-helper cells LSES. These SNPs map to *LINC0029*, a long non-coding RNA in proximity (+/-50kb) to *PRR5* and *ANP32BP2* (**Figure 5B**). Of the 10 SNPs with suggestive *p* values at this locus, three are in LD with the top SNP, rs11914148 (0.6 < r^2^ < 0.8), but the remaining SNPs, have weaker r^2^ values (r^2^ < 0.4), suggesting an independent secondary signal at this locus. The effects of the top SNP demonstrated some heterogeneity across cancer types, particularly for adrenocortical carcinoma (ACC), esophageal carcinoma (ESCA), and lung squamous cell carcinoma (LUSC) (**Figure 5C**). The locus on chr 8 associated with Tfh cell LSES represented by one genome-wide significant SNP, rs71510648, and five suggestive SNPs, rs11989637, rs35312266, rs13281971, rs13259019 and rs71510649, map to the *SH2D4A* gene and is in proximity to *CSGALNACT1* gene (+/-50kb) (**Figure 5D**). The effect of the top SNP within each tumor was largely consistent with the pan-cancer analysis except for mesothelioma (MESO) which displayed a strong trend in the opposite direction (**Figure 5E**).

Two more loci included SNPs with relatively low minor allele frequency (MAF; 0.005 < MAF 0.01). One SNP on chr 2, rs78868990, was strongly associated (p=9.66 x 10^-10^) with CD8 T-cells LSES and also has a suggestive association with Cytotoxic T-cell LSES. In addition, a second SNP, rs729684, shows suggestive association with CD8 T-cells LSES. These two SNPs map to *SPAG16* and are within 100kb downstream of *IKZF2*, which encodes for Ikaros family member hematopoietic-specific transcription factors (**Figure S5A**). In addition, we found four SNPs in chr 17 with multiple genome-wide significant and suggestive associations with three immune traits associated with T-cell subsets: rs112236917, rs112262673, rs73316909 and rs138156694 were significantly associated with T-helper cells LSES; rs112236917, rs73316909 and rs138156694 had significant associations and rs112262673 had a suggestive association with CD8 T-cells LSES; and rs112236917, rs73316909 and rs138156694 had suggestive associations with Tfh cells LSES. All four SNPs map to *TRPV3*, and are in proximity to *SPATA22*, *ASPA* and *TRPV1* (**Figure S5C**). These effects across tumor types for these rare variants demonstrated no significant heterogeneity but the effect estimates were a relatively less consistent across cancer subtypes, likely due to statistical fluctuations due to their low allele frequencies (**Figure S5B, and S5D**). Finally, we found a suggestive association between CD8 T-cells LSES and rs4495109 at the *KDR* locus (**Figure S5E**), a gene which has previously been associated with immune infiltration in colon cancer (Van den Eynde et al., 2018). However, the SNP associated with immune microenvironment effects in that report, rs1870377, was not significantly associated (*p* = 0.12) with CD8 T-cells LSES in our analysis. Of note, the effect of rs4495109 on CD8 T-cells was inflated in colon adenocarcinomas (COAD), cholangiocarcinomas (CHOL) and endometrial carcinomas (UCEC) relative to the pan-cancer analysis (**Figure S5F**).

### Association with high penetrance susceptibility variants

We performed association analyses between pathogenic and likely pathogenic germline mutations (referred here as rare variants) in both protein coding regions and high penetrance susceptibility genes, immune traits and immune subtypes (**Figure 6**). Since mutations in most of the genes were rare, we collapsed, when possible, genes into categories summarizing different biologic processes or functions (**Figure S6A**). However, as mutations within the homologous recombination repair (HR) genes *BRCA1* and *BRCA2* were more common, we analyzed mutations within each of the genes separately. Overall, 21 genotypic variables were used (**Figure S6B**).

**Figure 6.**
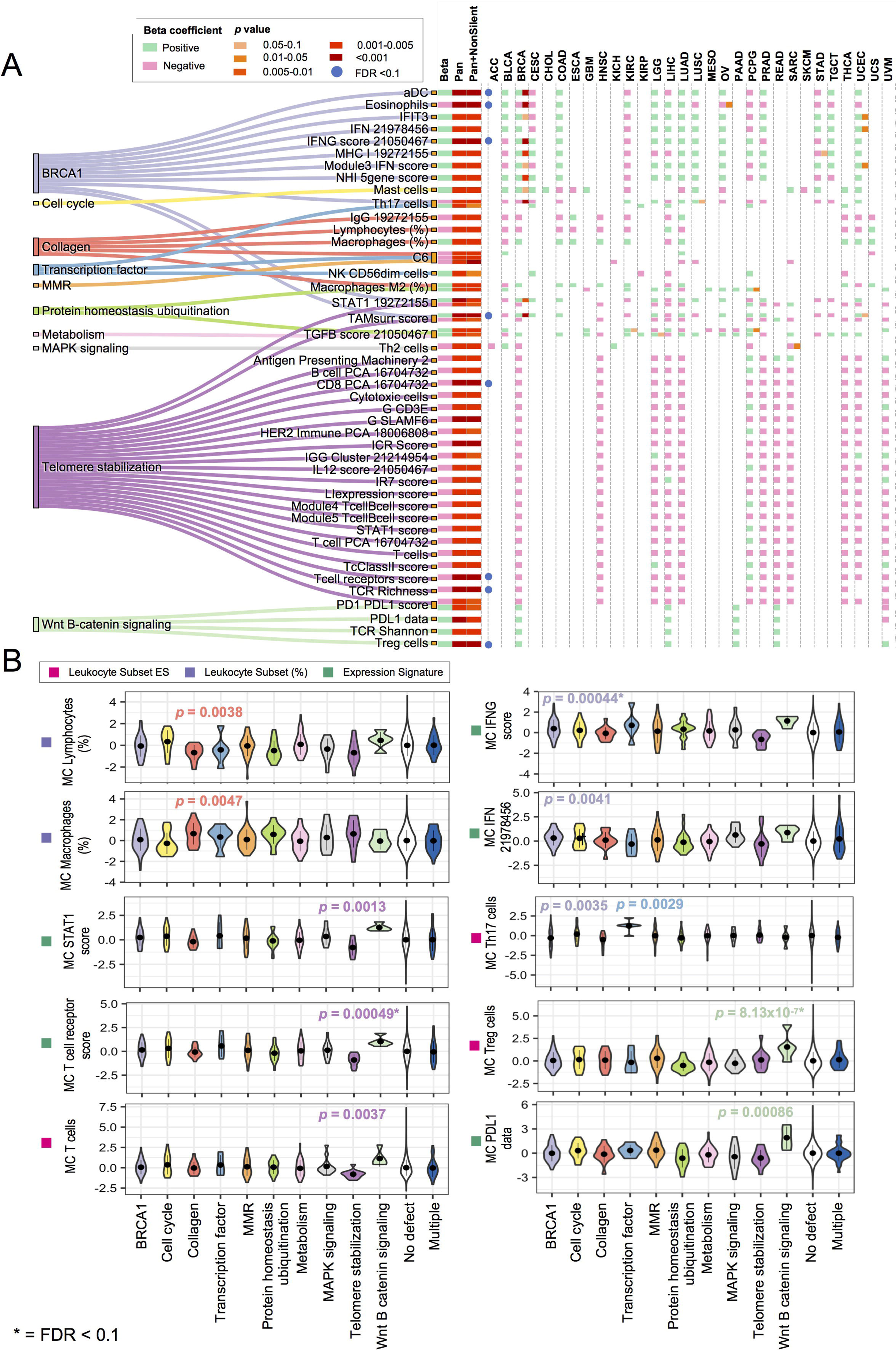
Pathogenic or likely pathogenic variants modulating immune traits. (**A**) Suggestive associations (*p* < 0.005 and FDR *p* ≤ 0.25) between rare genetic pathogenic or likely pathogenic variants extracted from whole-exome data (Huang KL, et al., *Cell*, 2018) grouped by curated mutually exclusive functional categories (*left nodes*) and immune traits (*right nodes*) as identified in pan-cancer regression models adjusted for cancer type, age, sex, and PC1-7. Significant associations (*p* < 0.005 and FDR *p* < 0.1) are highlighted with blue dots. Within cancer, association tests are run with age, sex and PC1-7 as covariates, except in CESC, OV, PRAD, TGCT, UCEC and UCS where only age and PC1-7 are used. The adjusted-*p* for the non-silent mutation rate are also shown. Germline genotypic variables with a number of events lower than five across cancers were excluded from the analysis. Beta coefficients and significance level are visualized pan-cancer and per cancer (*right side*). Significance per cancer was only calculated when number of events was greater than two. The Beta coefficient is shown irrespectively of the significance and number of events. Beta: beta coefficient. Pan: *p* value pan-cancer. Pan+Nonsilent: *p* value pan-cancer adjusted for non-silent mutation rate. **(B)** Values of representative immune traits (mean centered by cancer type) are displayed across samples with mutations in genes related to the defined functional categories.

In an analysis across all cancer types, we found significant associations (FDR *p* < 0.1) between at least one immune trait and mutations in *BRCA1* genes, WNT-Beta catenin, and telomere-stabilization pathways. In addition, we found suggestive associations (*p* < 0.005, FDR from *p* > 0.1 to *p* ≤ 0.25) between at least one immune trait and six other categories: cell cycle, collagen, transcription factor, mismatch repair (MMR), protein homeostasis ubiquitination, metabolism, and MAPK signaling (**Figure 6A-B**). These associations were only minimally altered by correction for the somatic mutational load (**Figure 6A**). We also tested the relation between rare variants and parameters reflecting somatic DNA alterations (**Figure S6C-D**). However, since germline mutations in MMR genes are known to affect the somatic mutation rate, which is an important predictor of immune responsiveness, we performed more detailed analyses of the effects of these mutations.

As expected, MMR germline variants were associated with a higher mutational/neoantigen load and a higher microsatellite instability score (MANTIS) (Middha et al., 2017) (**Figure S6C**). These associations were statistically significant only in COAD and UCEC cancers. MMR germline mutations were associated with higher leukocyte infiltration only in colon cancer (**Figure 7A**). Overall, a higher leukocyte fraction in MMR germline mutated samples were confined to the ones acquiring the microsatellite instability phenotype (**Figure 7B**). Interestingly, among microsatellite unstable tumors (MSI), the ones driven by MMR germline mutations had higher leukocyte infiltration (**Figure 7B**). A similar trend was observed for other immune signatures (data not shown).

**Figure 7.**
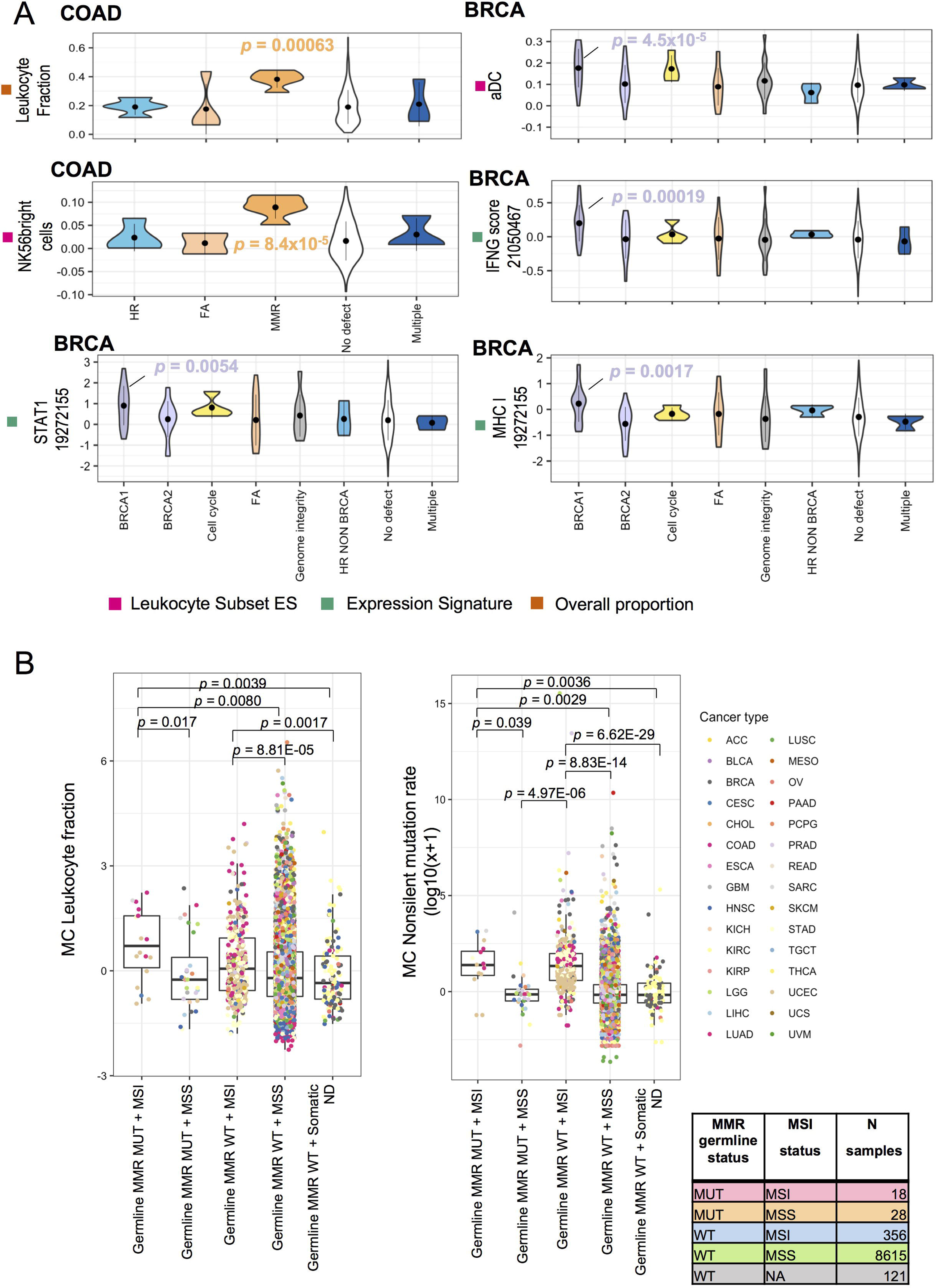
Representative immune traits modulated by pathogenic or likely pathogenic variants in breast and colon cancer and according with microsatellite instability status. **(A)** Representative associations between pathogenic or likely pathogenic variants, grouped by functional categories, and representative immune traits within colon adenocarcinoma (COAD) and breast invasive carcinoma (BRCA). **(B)** Leukocyte fraction and non-silent mutation rate (mean centered by cancer type, (MC)) by combined germline mutation status across mismatch repair (MMR) genes and somatic microsatellite instability (MSI) status as identified by MANTIS score (threshold = 0.4, Middha et al., 2017). Five categories are displayed: MMR germline mutated (MUT) and microsatellite unstable (MSI, n=18), MMR germline MUT and microsatellite stable (MSS) (n=28), MMR germline wild type (WT) and MSI (n=356), MMR germline WT and MSS (n=8615), and MMR germline WT and MSI MANTIS not determined (ND) (n=121). Samples are colored by cancer type. *P* values < 0.05 from regression analyses adjusted for sex, age, cancer type (for pancancer analyses, (B)), and PC1-7, are displayed. FA: Fanconi Anemia; HR NON BRCA: homologous recombination excluding *BRCA1* and *BRCA2*; MMR: mismatch repair; NER: nucleotide excision repair.

Germline mutations in Fanconi Anemia (FA) and in *BRCA1/2* genes were strongly associated with a higher Homologous Recombination (HR) defect score (**Figure S6B**). However, only mutations of *BRCA1* were associated with favorable immunologic parameters such as the higher values of MHC and traits that are part of the IFN module and the abundance of activated dendritic cells (aDC) which is also part of the IFN module. These associations were driven by breast cancer samples (**Figure 6A and Figure 7A**). In addition, when the analysis within breast cancer samples was corrected by intrinsic molecular subtypes (i.e. basal vs. luminal vs. HER2-enriched), the associations were no longer significant.

Finally, mutations in genes involved in telomere stabilization were strongly correlated with features associated with T-cell exclusion, including low abundance of T-cell and cytotoxic cells and diminished IFN-related signatures (including STAT1), and TCR richness. Germline mutations of a collagen-related related gene (*COL7A1*), were nominally significantly associated with increased macrophage infiltration accompanied by decreased lymphocyte infiltration. In addition, the WNT-Beta catenin germline mutations were consistent with the presence of an inflammatory phenotype accompanied by counter-regulatory mechanisms such as the activation of the PD-L1 signaling and the recruitment of T-regulatory (T-reg) cells.

## Discussion

We evaluated the effect of germline variation on immune response in tumors using genome-wide heritability, genome-wide association for common variants, and gene-based analyses focusing on rare variants in known cancer susceptibility genes. We identified significant genome wide heritability in pan-cancer analyses of 33 immune traits, particularly traits representing T-cell enrichment and activation of Interferon-gamma. We also identified 23 loci that pass genome-wide significance for association with heritable immune signatures. Finally, our rare variant analyses demonstrated expected associations between mutations in MMR genes and immune signatures in colon cancer, as well as previously unreported associations between mutations in genes in several pathways and cancer immune signatures. Thus, overall, we identified a complex set of germline effects on tumor immune response.

Pan-cancer common variant heritability analyses identified significant heritability for several of the tumor immune response traits. The most heritable immune traits were members of the Cytotoxic module, which were developed based on genes exclusively expressed in those cell subsets. These traits were originally developed to identify subsets of immune cells infiltrating tumors and are strongly associated with survival. They were all highly correlated despite the traits being comprised of different genes. As expected, their heritability values were very similar, with a value of ∼20% heritability. Furthermore, top hits from GWAS frequently demonstrated correlations for these traits. The correlations in the genotypic results suggest that these traits are genetically similar and the cell types they represent are driven into the tumor by similar mechanisms. In addition, since the genes in the expression signatures for each of these cell types were selected to be distinct, the genetic correlation between these traits was unlikely due to traditional expression quantitative trait loci (eQTLs) and rather represented genetic effects on the cellular infiltrates and/or the activation states of these cells. The second most heritable group of immune traits were members of the interferon (IFN) module, which had ∼15% genome wide heritability. Traits that were members of this module were also highly correlated, had very similar heritability estimates, and had overlapping genome wide association loci, suggesting that their heritability estimates are robust to the gene subsets selected for the signature and represent an aspect of immune microenvironment.

The heritability that we identified were on par with many other complex traits in humans, such as body mass index and fasting glucose (Shi et al., 2016). Genome-wide heritability derived from array-based data is generally considered to capture only a fraction of the overall heritability of a trait for several reasons; most prominently, it misses rare variation (Zaitlen and Kraft, 2012). TCGA only includes common variant germline data and not whole genome germline sequencing. A recent study using whole genome sequencing to calculate heritability for height found that rare variants account for a large fraction of the heritability and recover all of the “missing heritability” previously calculated based on pedigree studies (Wainschtein et al., 2019). Thus, it is likely that full sequencing may also substantially increase the estimates of heritability of these immune signatures.

We also found significant interactions between heritability and tumor immune subtypes. In particular, some traits that showed pan-cancer heritability, such as members of the Cytotoxic module, also had significant interactions with tumor immune subtypes. In addition, some of the immune traits that were not significant over all cancer types, showed significant interactions, including those that capture TGF-β signaling and macrophage infiltration. These interactions suggest that, for some immune traits, the germline effects may be context dependent. One possibility is that the germline may be acting as a modifier of other factors that determine the tumor immune subtype. For most of the immune traits that interacted with immune subtypes, the heritabilities were higher in the wound healing quiet subtype, the immune subtype with lowest immune cell infiltrations of those analyzed. This pattern suggests that, for these traits, the main driver of the immune microenvironment in the most immune infiltrated tumors is not common genetic variation, but that the germline may be modifying the immune features in the tumors with the lowest level of immune infiltration.

Many immune traits did not have common variant heritability in either pan-cancer analyses or in interaction analyses. In particular, the degree of overall leukocyte infiltrate was not heritable. However, our analyses do not capture rare variant heritability. In fact, in one instance of rare pathogenic mutations in mismatch repair genes in colorectal cancer, we found that the leukocyte fraction was dramatically increased, consistent with the known effect of immunotherapy (Le et al., 2015, 2017a). In addition, some common variant effects may be cancer specific, and we were under-powered to detect them.

In genome wide association analyses, we found two significant loci for interferon-gamma signatures. One locus at chr 2 included 2 SNPs within the *IFIH1* gene, rs2111485 and rs1190760 in high LD. These SNPs have been associated with a variety of autoimmune disorders including type 1 diabetes mellitus (T1DM) (Smyth et al., 2006), vitiligo (Jin et al., 2012), psoriasis (Tsoi et al., 2012), inflammatory bowel disease (Liu et al., 2015), and systemic lupus erythematosus (Robinson et al., 2011), demonstrating a genetic link between autoimmunity and the immune response to cancer. *IFIH1* is induced by interferon and acts as an RNA-dependent ATPase (Kang et al., 2002; Yoneyama et al., 2005). One of these SNPs, rs1190760, is a non-synonymous variant (A946T) in *IFIH1* and may be the causal variant at this locus, although these SNPs are also associated with expression (Liu et al., 2009). The A allele of rs2111485 and C allele of rs1190760 were associated with increased risk of T1DM and vitiligo and were associated with increased IFN signaling signature in our dataset, suggesting a positive correlation between genetic risk for T1DM, vitiligo, and interferon signature in tumor. Since interferon signatures have been associated with a positive prognosis for patients on immunotherapy (Ayers et al., 2017), our results suggest that this SNP may also be associated with efficacy among patients receiving immunotherapy, which is consistent with preliminary data in melanoma patients receiving checkpoint inhibitors (Chat et al., 2019). In contrast, the A allele of rs2111485 and C allele of rs1190760 are associated with decreased risk of psoriasis and inflammatory bowel disease (Liu et al., 2015; Tsoi et al., 2012). Our analyses identified at least one additional suggestive (*p* < 10^-6^) association at this locus characterized by rs17716942, a SNP which has been associated with psoriasis (Tsoi et al., 2012).

Our analyses also identified a tumor-immune associated cluster of SNPs on chr 3 near *TMEM108*. These SNPs were nominally associated with several other cellular signatures including one for activated dendritic cells and one for the fraction of NK cells. *TMEM108* is not known to have an effect on interferon signaling, but may signal through the WNT-Beta Catenin pathway (Jiao et al., 2017). *TMEM108* is also a cancer-testis (CT) antigen. CT antigens are typically only expressed in normal testis and in other developmentally regulated tissues (e.g. placenta), but are aberrantly expressed in many types of cancers (Scanlan et al., 2004). Due to their restricted expression pattern, CT antigens are frequently recognized by the immune system of cancer patients. Interestingly, 4 SNPs in chr 17 that had suggestive cross-associations with three immune traits associated with T cell subsets mapped in proximity to the *SPATA22* gene, which is also a CT antigen (da Silva et al., 2017; Wang et al., 2016).

We also found 11 genomic loci showing significant association with traits of T-cell subsets and/or NK cells, and some of these also showed genome-wide significance for signatures of other T-cell subsets. One locus on chr 22 had a genome-wide significant association with a signature for CD8 cells LSES and T central memory LSES cells, and a suggestive association with a signature for T helper cells LSES. The top SNPs at this locus were in a long non-coding RNA, linc00229. This RNA is upregulated in the context of T cell response to infection (Fu et al., 2017). Another set of SNPs on chr 8 that have a genome wide significant association with Tfh cell LSES map to the locus for the *SH2D4A* gene. This gene is required for T-cell antigen-induced cytokine synthesis in lymphocytes (Marti et al., 2006), suggesting that this locus may mediate its effect directly on T-cell function. Another locus on chr 2 that was associated with CD8 T cells was within ∼100 KB of *IKFZ2*, a major transcriptional regulator of lymphocyte differentiation (Georgopoulos et al., 1997). Our analyses also identified genome-wide significant SNPs at a locus within *TPRV3* and near *TPRV1*. *TRPV1* and *TRPV3* encode Transient Receptor Potential (TRP) ion channels of the Vanilloid (TRPV) subfamily. *TRPV3* is predominantly expressed on keratinocytes. *TRPV1* is the receptor for the phytochemical capsaicin, which has been shown to be expressed on cells of the immune system as well as several types of cancer cells. *TRPV1* is also activated by bacterial lipopolysaccharide and has been suggested to be a potential link between inflammation, cancer, and immunity (Bujak et al., 2019). In addition, we found a suggestive association between CD8 T-cell LSES and a SNP (rs4495109) in the *KDR* locus on chr 4. Interestingly, in colorectal cancer, a germline polymorphism in *KDR* Q472H (rs1870377) was strongly associated with the immunoscore (density of CD8 and CD3 cells in the invasive margin and center of the tumor) (Van den Eynde et al., 2018), although this SNP did not replicate in our analyses. The detection of different KDR SNPs might be the result of different approaches. For instance, the Immunoscore measures the spatial distribution of two distinct leukocyte subpopulations, while here we estimated the abundance of leukocyte populations based on expression data. Additionally, only colorectal cancer patients were analyzed in Van den Eynde et al. while we employed a pan-cancer approach. In addition, Van den Eynde, et al. used a targeted panel focused on coding sequence variants that did not include rs4495109, which was the SNP that was most strongly associated in our data. Despite these differences, the fact that both of these studies identify variants in KDR highlights the importance of this gene that functions as one of the receptors of VEGF, which has an immune suppressive effect (Voron et al., 2015).

Rare variants analyses demonstrated intriguing associations between genetic variants related to cancer development and intratumoral immune response. We found two associations between DNA-repair genes and immune traits. First, we found that *BRCA1* germline mutations were associated with higher levels of favorable immune parameters. These associations were not observed for *BRCA2*, FA, and non-BRCA homologous recombination defects, despite the comparable number of germline mutations in these genes. Moreover, these effects were restricted to breast cancer samples, and driven by the higher rate of basal-like/triple negative phenotype among *BRCA1* mutation carriers with breast cancer. Basal-like breast cancers have previously been shown to be more likely to have a more robust immune infiltration compared with other breast cancers (Jézéquel et al., 2015), and our analyses suggest that *BRCA1* may mediate its effect on immune response by its effect on breast cancer subtypes. We also found that mutations in genes in MMR pathway are associated with a more robust immune response and that this is likely mediated by the microsatellite instability phenotype. These results are consistent with previous reports and with the responsiveness to checkpoint inhibitors of patients carrying germline MMR mutations (Le et al., 2017b).

Patients with germline mutations of telomere-stabilization genes had lower lymphocytic infiltration. The increased risk of cancer development in patients with defects in genes related to telomere stabilization might be due to rearrangements through chr breakage-fusion-bridge cycles triggered by critically short telomeres (Aviv et al., 2017; Maser and DePinho, 2002). These events might lead to genomic instability. While a higher degree of aneuploidy has been associated with reduced lymphocytic infiltration in some tumors (Davoli et al., 2017), we did not detect any association between mutations in telomere-stabilization genes and measures of aneuploidy. Another potential mechanism through which these alterations might lead to lower lymphocytic infiltration could be related to the functional modification of lymphocyte proliferative capacity following antigen recognition. Shorter telomere length has been associated with lower persistence of adoptively transferred lymphocytes and decreased response to treatment (Rosenberg et al., 2011). Additionally, decreased leukocyte telomere length has been correlated with reduced survival and altered immune functions in patients with colorectal cancer (Chen et al., 2014).

The associations between germline mutations of genes in the WNT-Beta catenin pathways and increased levels of T-cell and counter-regulatory mechanisms (PDL1 and T-reg), supports the critical role of this pathway in modulating anti-tumor immunity. In fact, somatic alterations leading to WNT-Beta catenin pathway activations have been associated with T-cell excluded immune phenotypes in multiple tumors (Luke et al., 2019; Spranger et al., 2015). This is likely due to the decreased secretion of CCL4 following WNT-Beta catenin activation, resulting in decreased recruitment of BATF3 DC recruitment and subsequent lack of T-cell priming and chemoattraction (Spranger et al., 2017). While germline mutations of *APC* and *PTCH1* (the WNT-Beta catenin negative regulators with deleterious germline mutations in the present dataset) might predict pathway activation, it is possible that such alterations, when present in the germline, induce a pathway’s downregulation at the somatic level by triggering compensatory mechanisms.

One prior study identified many eQTLs in TCGA, some of which may be related to immune signatures (Lim et al., 2018). However, this study focused mostly on eQTLs, including eQTLs that may affect immune related genes. These investigators also performed analyses within cancer types and identified 63 SNPs with *p* values that passed their threshold of significance. However, all of these loci were associated within one cancer type and the majority were in the stomach adenocarcinoma (STAD) subtype. Our study is different in several important ways. First, we began with a more comprehensive list of 139 immune traits evaluated recently (Thorsson et al 2018) and calculated heritability using genome-wide common variant data. We then prioritized traits that showed evidence for heritability for GWAS. We performed GWAS using pan-cancer analyses which increased our power. Finally, we also evaluated the effects of rare variants in cancer susceptibility genes to build a more complete picture of the impact of germline variation on tumor immune response.

Our analyses had several limitations. While our heritability analyses were performed using >7000 samples and our GWAS analyses included >9000 samples, GWAS and heritability analyses for many complex traits often use larger sample sizes as the effect sizes are often small and the adjustment for multiple hypothesis testing is substantial. Thus, future studies in larger sample sizes are likely to identify more loci which affect the immune trait microenvironment. Furthermore, our analyses focused on detecting pan-cancer genome-wide heritabilities and loci. There may be tissue specific or other conditional factors that may affect either individual loci and/or genome-wide heritability. Despite this limitation, we did find many significant loci, suggesting that at least some germline variants affect the immune microenvironment of many different types of cancers. While we did search for some interactions using the discrete immune subtypes, there may be other subsets of cancers (e.g. adenocarcinoma vs. squamous cell, high vs. low neoantigen load) which also interact with germline factors. In addition, many of our findings were driven by signatures that were defined by gene expression. Since gene expression is often controlled by variants, some of the loci that we detect may not be controlling the overall level of immune response, but could be affecting the expression of the genes within the cells as expression quantitative trait loci (eQTLs). Although we cannot formally exclude this possibility, we believe this is relatively unlikely for most of the single variants we present here. First, for 21 of the 23 genome-wide significant loci that we identified, the SNPs were not in *cis* (<100 KB) with genes that were part of the immune trait. Second, in the 2 cases in which we did identify strong eQTLs in cis (*HLA* locus SNPs associated with MHC and *IL7RA* locus SNPs associated with Th17 LSES), the SNPs were exclusively associated with a single trait (**Figure 3B**). In contrast, for the other loci we identify, the top variants have effects on multiple immune traits, suggesting a more robust effect on the immune infiltrate rather than an eQTL.

Our study leveraged TCGA, which collected specimens prior to the large-scale use of immunotherapy in cancers and very limited immunotherapy treatment information is recorded in TCGA clinical annotations. Therefore, we cannot make any conclusions about the genome wide or locus specific findings and their effects on immunotherapy. Germline variants in mismatch repair genes are already considered a strong predictor of response to checkpoint inhibitors and are one of the FDA indications for treatment with these agents, regardless of tissue of origin. In addition, the common variants we identify may be candidates for both predictors of treatment efficacy and/or immune related adverse events.

In summary, we have identified evidence of both common germline variant effects on tumor immune response and rare mutations in susceptibility genes that may affect tumor immune response. These effects should spur future research to determine whether these germline variants affect response to existing immunotherapies. In addition, they may suggest other pathways that could yield new targets for immunotherapy.

## Acknowledgements

We are grateful to the Society for Immunotherapy of Cancer (SITC) for the logistical support of the investigator meeting within the SITC Cancer Immune Responsiveness Workshop (San Francisco CA, US, April 2018; Houston, TX, US, May 2019). We are also grateful to Noah Zaitlen and Andy Dahl for useful discussions on heritability interaction analyses. This work was funded in part by R01CA227466 and K24CA169004 to EZ, Qatar National Research Fund (QNRF) NPRP10-0126-170262 and NPRP11S-0121-180351 grants to DB, Associazione Italiana per la Ricerca sul cancro (AIRC) grant IG2018-21846 to MC, and the National Cancer Institute (NCI) grant T32CA221709 postdoctoral fellowship to RWS.

## Author Contributions

Conceptualization: V.T., D.W., F.M.M., E.Z., D.B. Design: R.W.S., M.S., V.T., Y.M., T.K., R.F.S., O.F.B., E.P.P., S.S., D.W., E.Z., D.B. Analysis, Computation, and Software: R.W.S., M.S., V.T., W.H., J.R., Y.M., F.F., R.F.S., O.F.B., D.H., S.H., N.S., E.Z., D.B. Interpretation: R.W.S., M.S., V.T., W.H., J.R., Y.M., F.F., T.K., R.F.S., O.F.B., E.P.P., M.Ca., C.S., D.H., S.H., R.E.G., N.S., L.R., M.L.D, S.S., D.W., F.M.M., J.G., M.Ce, E.Z., D.B. Writing-Review & Editing: R.W.S., M.S., V.T., W.H., J.R., Y.M., F.F., T.K., R.F.S., O.F.B., E.P.P., M.Ca., C.S., D.H., S.H., R.E,G., N.S., L.R., M.L.D, S.S., D.W., F.M.M., J.G., M.Ce, E.Z., D.B. Supervision & Funding Acquisition: E.Z., D.B.

## STAR Methods

## LEAD CONTACT AND MATERIALS AVAILABILITY

Further information and requests for resources should be directed to and will be fulfilled by the Lead Contact, Rosalyn Sayaman. This study did not generate new unique reagents.

## EXPERIMENTAL MODEL AND SUBJECT DETAILS

### Human Subjects

A total of 11,521 genotype files from participants across 33 different cancer types included in TCGA were downloaded (Downloaded May 30, 2018 from https://portal.gdc.cancer.gov/legacy-archive). The TCGA dataset has previously been described(Liu et al., 2018). We excluded all participants with hematological malignancies (diffuse large B-cell lymphoma, and acute myeloid leukemia) and thymoma since these could not be characterized for immune cell infiltration based on gene expression analyses. We included all participants who had genotype data from Affymetrix array on at least one normal sample (peripheral blood or matched normal tissue). After data cleaning (see below), the dataset for imputation included 10,128 individuals. Further removal of samples based genetic relatedness, availability of immune traits and covariate data resulted in a dataset of 9,603 individuals. Of these individuals, there was a total of 4,585 men and 5,018 women based on self-reported sex. The participants’ age ranged from 11yr to 90y with a median age of 61yr. Self-reported race and ethnicity were available from 8,510 samples, of whom 7,073, 825, 583, 20, and 9 samples were reported to be White/Caucasian, Black/African American, Asian, American Indian or Alaska Native, and Native Hawaiian/Pacific Islander, respectively. Self-reported ethnicity was reported on 7,351 samples, of whom 295 and 7056 samples were reported to be Hispanic/Latino, or Not Hispanic/Latino, respectively. Of these samples, 8,204 were typed on blood-derived normal, 1,397 on solid tissue normal and 2 on buccal cell normal tissue. Institutional review boards at each of the sites that provided samples and data reviewed the consent forms and approved the use of samples.

### Germline genotype data

Germline genotype data for common variants used in heritability analysis and Genome-Wide Association Studies (GWAS) were obtained from Affymetrix Genome Wide SNP 6.0 arrays (TCGA legacy archive https://portal.gdc.cancer.gov/legacy-archive). Birdseed genotyping files representing 905,600 variants for 11,521 samples were downloaded.

### Whole exome sequencing data

Germline genotype data for rare variants were based on whole exome sequencing data (TCGA archive https://portal.gdc.cancer.gov/). Processed whole exome sequencing data for pathogenic or likely-pathogenic variants from Huang et al., 2018 were considered for rare variant analysis representing 10,389 individuals of which 9,138 had all of the phenotype and covariate information for analysis.

### Immune traits

Immune traits considered for analysis were merged from two sources in (Thorsson et al., 2018): the Feature Matrix (56 immune related features selected) and the scores for 160 genes signatures in tumor samples (160 features, Scores_160_Signatures.tsv (GDC manuscript publication page https://gdc.cancer.gov/about-data/publications/panimmune).

## METHOD DETAILS

### Affymetrix Genome-Wide SNP 6.0 Quality Control

Birdseed files were read in R v3.5.0 using the Affymetrix SNP Array 6.0 (release 35) annotation file, and 905,422 variants were successfully loaded and analyzed in PLINK version 1.9. Samples were cross-referenced against previously whitelisted genotyping samples (Thorsson et al., 2018)]. Based on established TCGA barcode identifiers, samples annotated with Analyte code “G” (Whole Genome Amplification) were further excluded. A final set of 10,946 whitelisted samples with Analyte code “D” (DNA) were retained for quality assessment.

Stringent quality control measures were applied to the SNP genotyping data (Figure 1, top QC panel). SNPs and individuals with greater than 5% missingness were excluded; leaving a total of 861,351 variants and 10,917 samples for subsequent analysis.

Initial Principal Component Analysis (PCA) ancestry analysis was performed to facilitate heterozygosity calculations. PCA without linkage-disequilibrium pruning was performed in PLINK 1.9 (Chang et al., 2015), and visual examination of the concordance of the principal component plots with the self-reported race and ethnicity annotations revealed that the first 3-4 PCs captured population structure information, while PCs 5-6 captured outliers. PCA initial ancestry clusters were determined by performing both k-means and partition around medoids (PAM) clustering on either the first three or first four PCs. We computed gap statistics and average silhouette widths iteratively for number of clusters, k=1 to 10 for k-means and PAM methods respectively to find the optimal number of clusters for each method. We found PAM using the three PCs yielding 4 optimal clusters to show high concordance with self-reported race/ethnicity (ancestry cluster 1 = European, cluster 2 = Asian, cluster 3 = African, cluster 4 = American). Based on the initial ancestry cluster assignments, heterozygosity was calculated in PLINK 1.9 within each initial PCA-based ancestry cluster and a total of 250 samples with heterozygosity >3*SD above the ancestry mean were removed.

Selection of a representative sample for each individual was then conducted. Individuals represented by more than one sample, blood-derived normal samples were preferentially selected; for those with more than one blood-derived samples, samples with higher call rates were retained. After these steps, a total of 10,128 unique individuals remained for subsequent analysis.

Final filtering steps for SNPs were conducted across the 10,128 unique individuals and restricted to autosomal chrs. Hardy-Weinberg Equilibrium (HWE) was calculated in PLINK 1.9 across individuals within largest ancestry cluster (European ancestry cluster 1). SNPs that deviated from the expectation under HWE (*p* < 1×10^−6^) within the European ancestry cluster were excluded with the exception of SNPs previously associated with any cancer as reported in the GWAS catalog (*p* < 5×10^−8^) (Rashkin et al., 2019) since they may deviate from HWE in cancer patients. Minor allele frequency (MAF) was calculated and variants with MAF < 0.005 were excluded. Finally, duplicate SNPs with identical genomic first position were removed. A total of 838,948 autosomal chr variants for 10,128 unique individuals passed after the aforementioned QC steps.

### Stranding and Reference Panel Imputation

The quality-controlled genotyping file was stranded and imputed against the Haplotype Reference Consortium (HRC) (Loh et al., 2016a; McCarthy et al., 2016). Prior to HRC stranding, all palindromic SNPs (A/T or G/C) were removed. Stranding was then performed using the McCarthy Group tools (HRC-1000G-check-bim-v4.29), which compares our data genotyping alleles to the corresponding SNP alleles from HRC (v1.1 HRC.r1-1.GRCh37.wgs.mac5.sites.tab), leaving 680,389 correctly matched variants for imputation.

Phasing and imputation were performed using a standard pipeline on the Michigan Imputation Server (MIS). Phasing was performed using Eagle version v2.3 (Loh et al., 2016b) on the variant call file (VCF). To reduce the run time, the VCF file was divided into 22 files corresponding to individual autosomal chrs. By default, Eagle restricts analysis to bi-allelic variants that exist in both the target and reference data. Minimac3 was used to run the imputation. For each of the 22 VCF files, the MIS breaks the dataset into non-overlapping chunks prior to imputation. For HRC imputation, the HRC r1.1.2016 reference panel was selected using mixed population for QC, with a total of 39,127,678 SNPs returned after imputation.

### Final Ancestry Calls

PCA was performed on the final quality-controlled genotyping file and final PAM-based ancestry clusters were computed for the 10,128 individuals for optimal k=4. We found very high concordance of initial and final ancestry assignments (99.98% matching, the 2 samples varying between initial and final ancestry cluster computation assigned to NA).

The four ancestry cluster are as follows: (1) PAM ancestry cluster 1 is concordant with European ancestry, capturing 97.27% of individuals self-reporting as White, as well as 82.16% of individuals with self-reported non-Hispanic/non-Latino ancestry and 45.96% with self-reported Hispanic/Latino ancestry; (2) ancestry cluster 2 with African ancestry, capturing of 97.53% of individuals self-reporting as Black/African-American race; (3) ancestry cluster 3 with Asian ancestry, capturing 90.88% of individuals self-reporting as Asian and 88.89% self-reporting as Native Hawaiian/Pacific Islander; and (4) ancestry cluster 4 with a subgroup of individuals with American ancestry capturing 60% of individuals self-reporting as American Indian /Alaska Native and 47.2% with self-reported Hispanic/Latino ethnicity (Carrot-Zhang J et al, manuscript submitted).

PC’s 1-7 show further population sub-structure in the Asian and European ancestry clusters. PAM ancestry sub-clusters were computed using PC’s 1-7 for individuals within the Asian ancestry cluster which yielded two optimal sub-clusters, and within the European ancestry cluster which yielded three optimal sub-clusters (GDC Publication Page Figure S2-B). Of note, 72.46% of European sub-cluster 3 self-reports as Asian (15.94% have no race reported). Ancestry clusters, sub-clusters, self-reported race and ethnicity and PC’s 1-7 are provided for each individual.

### Feature Selection for Analysis

Immune traits considered for analysis were merged from two sources in (Thorsson et al., 2018): the Feature Matrix (56 immune related features selected) and the scores for 160 genes signatures in tumor samples (160 features, Scores_160_Signatures.tsv on GDC manuscript publication page https://gdc.cancer.gov/about-data/publications/panimmune)

The 216 features were then filtered at three levels: (i) Level 1 filtering removed redundant features based on overlap of the feature matrix and the 160 signature feature set; (ii) Level 2 filtering removed features with limited interpretability; (iii) Level 3 filtering removed features with highly skewed distributions, which would not be amenable to subsequent analyses. A final set of 139 features were used in subsequent germline analysis. These selected immune phenotypes encompass six broadly-defined immune trait phenotype categories: (1) Leukocyte Subset Enrichment Score (ES), which include 24 immune cell-specific activation scores as captured by single-sample gene set enrichment analysis (ssGSEA) (Bindea et al., 2013); (2) Leukocyte Subset Percentages (%), which include 268 immune cell relative proportion measures (22 individual cells and 4 aggregates, as estimated by CIBERSORT) Cibersort and four aggregates values) (Gentles et al., 2015; Thorsson et al., 2018); (3) Overall Proportion, which includes three measures, namely leukocyte fraction, stromal fraction and tumor infiltrating leukocyte (TIL) regional fraction; (4) Adaptive Receptor, which includes four scores related to T-cell and B-cell receptor (TCR and BCR) Shannon diversity and richness; (5) Expression Signature, which includes four ssGSEA scores specific to lymphatic vessels (Bindea et al., 2013), antigen-presenting machinery (APM1 and APM2)(Şenbabaoğlu et al., 2016), and angiogenesis (Şenbabaoğlu et al., 2016), a collection of 68 gene signatures related to immunomodulatory signaling including coherent modules specific to interferon signaling (IFN-γ), TGF-β, wound healing, (core serum response) and T/B-cell response catalogued from earlier studies (Amara et al., 2017; Wolf et al., 2014), and an Immunologic Constant of Rejection (ICR) signature summarizing a T-helper 1 (Th1)/cytotoxic polarization of the tumor microenvironment associated with favorable prognosis and responsiveness to immunotherapy (representing the functional orientation of the immune contexture (Bedognetti et al., 2016; Galon et al., 2013; Hendrickx et al., 2017); and (6) Attractor Metagene, which includes nine TCGA-based co-expression signatures (metagene attractors) (Cheng et al., 2013a, 2013b).

### Covariate selection

For each analysis we included age, sex, cancer types and genetic ancestry from the principal components (principal components 1-7) as covariates. Self-reported age (age at diagnosis in years) from clinical data and PC 1-7 were used as continuous covariates. Cancer type based on TCGA study assignment and curated genotype-imputed sex assignments were used as categorical covariates. Due to missing self-reported sex, and discrepancy in self-reported and genotype-type based sex in the data, we carefully curated sex assignments. We recovered sex information by using X chr homozygosity estimate (XHE) after removal of the pseudo-autosomal region of the X chr. For those with missing sex information, individuals with XHE < 0.2 were assigned as females and individuals with XHE > 0.8 were assigned as males (N=21 males, N=28 females). Self-reported sex assignments were curated with individuals self-reporting as males with XHE < 0.2 reassigned as female (N=20 reassigned as females), and individuals self-reporting as females with XHE > 0.8 reassigned as males (N=6 reassigned as males). For brevity in the remainder of the manuscript, we refer to curated genotype-imputed sex assignments simply as sex. For a subset of analyses, we also included immune subtype as a covariate, as indicated in the text.

## QUANTIFICATION AND STATISTICAL ANALYSIS

### Immune Trait Correlations and Clustering

We calculated Pearson’s correlation coefficients across all pairs of the 139 immune traits. We then used hierarchical clustering (complete agglomerative method in R) using 1-correlation as distance metric) to identify modules of the immune traits.

### Immune Trait Normalization for Heritability, GWAS and Rare Variant Analysis

For immune phenotypes that appeared approximately normally distributed or normally distributed after correction for immune subtype, we calculated heritability using the immune phenotype without any transformation/normalization. For immune phenotypes that were highly skewed we applied a log10 transformation, and those that were approximately normally distributed after transformation were analyzed as log-transformed values. For immune phenotypes that could not be normalized with log10 transformation (usually due to a large number of 0 values), we dichotomized them at the median (i.e., higher and lower than median). These phenotypes were treated as binary variables in subsequent analysis.

### Germline Analysis

To examine the contribution of germline genetic variation to the functional orientation of the immune microenvironment, we conducted three types of analyses: (1) parallel heritability analysis (N_EUR_=7,813, N_AFR_=863, N_ASIAN_=570, N_AMR_=209 individuals), (2) genome-wide association studies (GWAS) (N=9,603), and (3) rare variant analysis (N=9,138) across 30 different non-hematological cancer types in The Cancer Genome Atlas (TCGA) (Figure 1, middle Analysis panel). Number of individuals, N, for each analysis type represent the maximum number of samples, however analyses per immune trait may proceed using fewer samples if NA values for specific individuals are present in a given immune trait. All germline analysis described below were adjusted for cancer type, age at diagnosis, sex, and principal components 1-7 derived from PCA of genetic ancestry.

### Heritability Analysis

Estimates of genome-wide heritability of the 139 described immune traits were calculated using the genomic-relatedness-based restricted maximum-likelihood (GREML) approach implemented in GCTA 1.91.2beta (Genome-wide Complex Trait Analysis), which simultaneously models the effect of all genetic variants (MAF > 0.01) (Yang et al., 2010, 2011). GREML calculates a genetic relatedness matrix (GRM) as a measure of the genetic similarity of unrelated individuals (GRM < 0.05) and compares it to the similarity of the measured immunological traits to calculate the total narrow-sense contribution of genotypic variance to overall phenotypic variance, V(Genotype)/V(Phenotype) [(Yang et al., 2010, 2011). All GREML analysis use the default average information (AI) algorithm to run REML iterations.

Since calculation of the GRM results in biased relatedness estimates for pairs of individuals who have different ancestry, we restricted heritability analysis primarily within the European ancestry group (N_EUR_=7,813, M=701,189 SNPs, K=32,292,666 GRM elements), which constitutes the largest ancestry population; secondary analysis is performed in the smaller African (N_AFR_=863, M=791,989, K=409,060), Asian (N_ASIAN_=570, M=649,768, K=183,315), and American (N_AMR_=209, M=751,507, K=24,753) ancestry groups. The four genetic ancestry groups were derived from optimal partition around medoids (PAM) clustering (**Figure S2A**) of individuals in principal components 1-3 based on their genotyping data (**Figure S2B**).

Heritability analyses are generally only well-powered for sample sizes of >1000; therefore, only the European ancestry subgroup was adequately powered and were presented in the main results. However, since we have previously seen associations between several of these immune parameters with various ancestral populations (Thorsson et al., 2018), we performed the heritability analyses separately in African, Asian, and American ancestral populations and present these in the supplementary results. To reduce bias in the heritability estimates, we removed one of each pairs of related individuals with A_jk_ > 0.05 (calculated from SNPs with MAF > 0.01) prior to running GREML. We calculated heritability using an unconstrained approach (allowing heritability to be < 0). Constraining the heritability to a range of 0-1 may lead to an upwards bias of the low heritability values, which is likely to be worse in smaller datasets. We used the likelihood ratio test (LRT) implemented in GREML to test if heritability is different than zero for each of the immune traits analyzed and used Benjamini-Hochberg false discovery rate (Benjamini and Hochberg, 1995) to calculate the false discovery rate (FDR). We present both FDR adjusted *p* values and unadjusted *p* values in the manuscript.

We also used GREML population to determine whether there are any contextual factors that interact with genome-wide common variant effects, including the major immune subtypes as determined by Thorsson et al and somatic mutations (divided into tertiles and dichotomized at 10 MB). We implemented the gene x environment (GxE) feature calculation in the European in GREML. For those immune traits for which we found nominally significant (*p* < 0.05) interactions, we calculated heritability in each stratified subset. For GxE calculations, the LRT tests the significance of the variance of GxE interaction effects.

### Genome-Wide Association Studies (GWAS)

We selected each of the immune phenotypes that demonstrated nominally significant genome-wide heritability (N=33) for GWAS. GWAS was conducted on all of the genotyped SNPs that passed QC and all of the imputed SNPs that had imputation R^2^ > 0.5 and minor allele frequency > 0.005 in the 9,603 unrelated individuals (PLINK 1.9 identity by descent, IBD, pihat < 0.25). Minor allele frequencies were recalculated post-imputation for only the subset of 9,603 individuals (PLINK 1.9). Of the 39,127,678 SNPs available after imputation, 10,955,441 passed both imputation quality and frequency thresholds and thus were included in the association analysis.

GWAS was performed using PLINK 1.9. Immune phenotypes that were approximately normal or normal after stratification by covariates were tested for association with SNPS using linear regression with covariates as above. Immune traits that were dichotomized for heritability analyses, were analyzed using logistic regression models. For each GWAS we also calculated the genome-wide inflation coefficient (lambda). We used the traditional cutoff of *p* < 5×10^−8^ as a cutoff for genome-wide significance and *p* < 1×10^−6^ to denote suggestive loci. Since we only selected the subset of phenotypes that was heritable and since many of the phenotypes are highly correlated, we did not correct the GWAS for the number of phenotypes analyzed. SNPs are annotated based on spanned genomic ranges (R v3.5.0, Bioconductor package GenomicRanges_1.32.6) with rsIDs (R v3.5.0, R package snplist_0.18.1, Bioconductor package SNPlocs.Hsapiens.dbSNP144.GRCh37_0.99.20) and with genes within +/-50kb of the SNP (R v3.5.0, Bioconductor package biomaRt_2.36.1 using grch37.ensembl.org as host). All annotations are based on Homo sapiens (human) genome assembly GRCh37 (hg19) from Genome Reference Consortium.

### Rare Variant Analyses

For rare variant analysis, we focused on well-annotated, pathogenic or likely pathogenic germline variants as previously defined (Huang et al., 2018). Exome files related to samples for which all the covariates (age, imputed sex, PC 1-7, and cancer type) and at least one immune trait was available were retained (N = 9,138). There were 832 pathogenic/likely pathogenic SNPs/Indels events with at least one copy of rare allele in the whole exome sequencing data, corresponding to 586 distinct pathogenic SNPs/Indels mapping to 99 genes.

We performed a pathway burden analysis using selected pre-defined biological pathways such as DNA damage repair and oncogenic processes, pan-cancer and per cancer (Huang et al, 2018). These pathways were manually curated to generate a list of mutually exclusive pathways. The only genes that were not collapsed into pathways were *BRCA1* and *BRCA2* for which a sufficient number of events across cancers exist. In the per-cancer analysis, we only included genes or pathways with at a number of events (mutations) greater than 4, including a total of 90 genes. In the per-cancer analysis, we only included genes or pathways with at least 3 events in the analyses. For each pathway, variants that fall within its selected set of genes were collapsed based on the presence or absence of any rare variant (i.e., 0 if no rare variant was present and 1 if there is at least one variant). We conducted a regression analysis to assess the association between the pathways’ burden of rare variants and immune traits. Traits assessed in this analysis are the same as the ones used for heritability analyses, with the addition of the immune subtypes (C1, C2, etc.), DNA-alteration related metrics such as the mutational load, the neoantigen load, the degree of copy number alterations (Thorsson et al., 2019), and the microsatellite instability score (MANTIS) (Middha et al., 2017). All regression models included the following covariates: age, imputed sex, PC 1-7. An additional covariate, cancer type, was included in the pan-cancer analysis. To check whether the results were driven by the mutational load, we ran regression models that include this variable as a covariate in the regression model.

In the pan-cancer analysis, we used a Benjamini-Hochberg false discovery rate (Benjamini and Hochberg, 1995) to correct for multiple hypothesis testing, accounting for all 21 genes and pathways tested and 154 phenotypes (139 immune traits, 9 DNA related metrics, and 6 immune subtypes). We used a cutoff of FDR *p* < 0.1 to identify significant gene/pathway-immune trait associations and a threshold of nominal *p* < 0.005 (FDR *p* ≤ 0.25) to identify suggestive associations. We used a more permissive cut-off in these analyses than the ones used in the heritability and GWAS to reduce type II error due to the low number of events (germline mutations).

## DATA AND CODE AVAILABILITY

The TCGA birdseed genotyping data and clinical data can be found at the legacy archive of the GDC (https://portal.gdc.cancer.gov/legacy-archive). The Cancer Genome Atlas (TCGA) quality controlled Genome Wide SNP 6.0 genotyping data imputed to Haplotype Reference Consortium are controlled access files, and will be made accessible to researchers with proper dbGAP authorization. These data have been deposited to Synapse (Available upon acceptance for publication). Details for software availability are in the Key Resources Table. The code generated during this study has been deposited to github (Available upon acceptance for publication)].

## Supplemental Figure Titles and Legends

**Introductory Supplementary Figure related to Figure 1.**
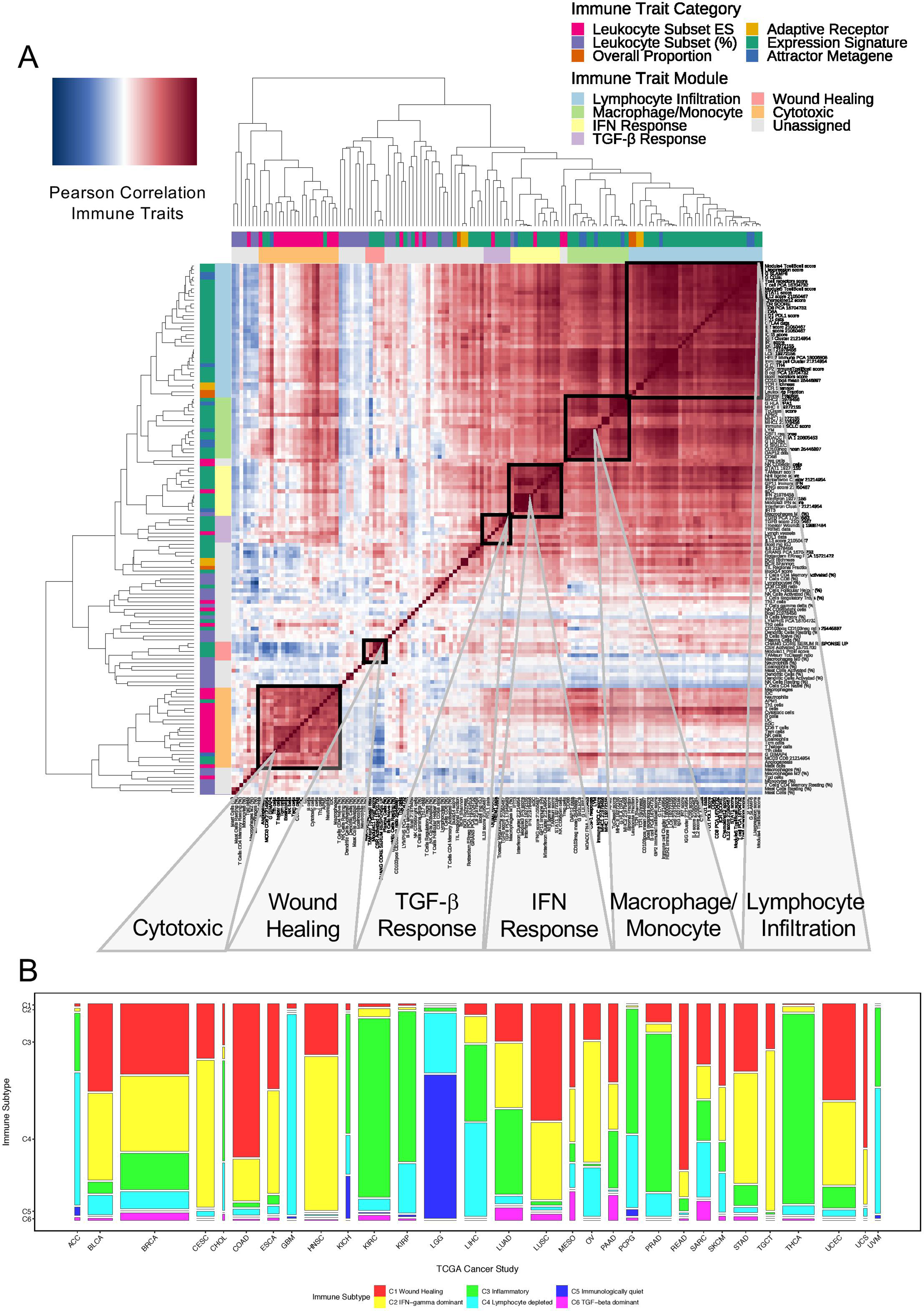
Correlation of immune traits and distribution of immune subtypes. One hundred thirty-nine immune traits encompassing measures of leukocyte subset enrichment score (ES), percentages (%), as well as overall proportion of leukocyte, stromal and TIL regional fraction, adaptive receptor BCR/TCR richness and Shannon diversity, immunomodulatory expression signatures and attractor metagenes, were considered for analysis. **(A)** Pairwise Pearson correlation matrix of the 139 immune traits show blocks of highly correlated immune traits which we define as modules: (i) lymphocyte infiltration, (ii) macrophage/monocyte, (iii) IFN-response, (iv) TGF-β response, (v) wound healing, and (vi) cytotoxic). Immune trait modules and corresponding immune trait categories are annotated by the color bar. Previous analysis of the distribution of these correlated immune signatures (immune modules) across the TCGA cohort led to identification of six distinct immunological subtypes across multiple cancer types: C1 Wound Healing, C2 IFN-γ dominant, C3 Inflammatory, C4 Lymphocyte depleted, C5 Immunologically quiet, and C6 TGF-β dominant (Thorsson V, et al., *Immunity*, 2018). **(B)** Mosaic plot showing a fractional representation of the number of individuals within each immune subtype across the 30 non-hematological cancers for the 9,603 TCGA subjects with available genotyping data used in the germline analysis of immune traits.

**Supplementary Figure related to Figure 2.**
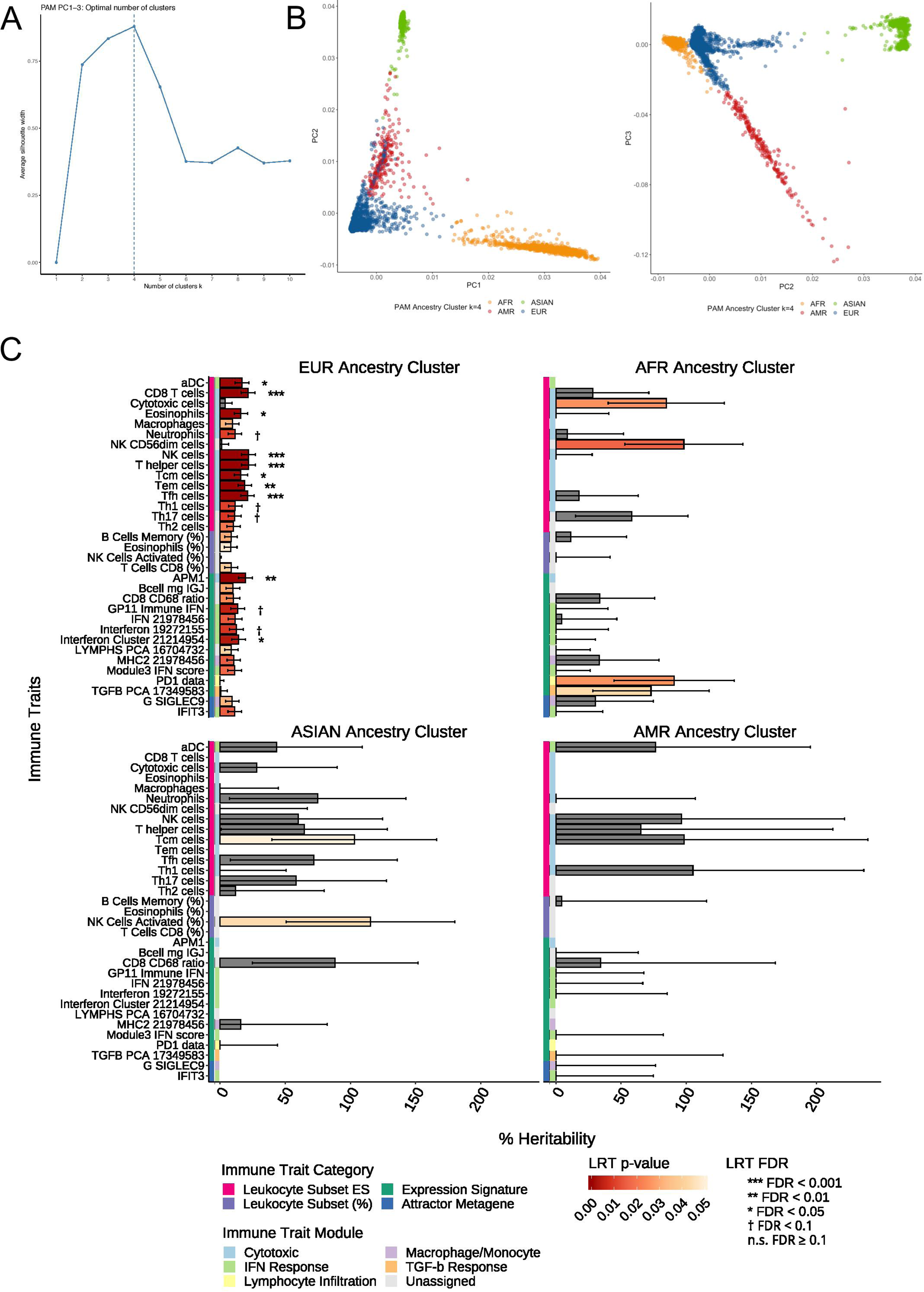
Genome-wide heritability of immune traits with ancestry groups. To calculate the pairwise relatedness of individuals in the genetic relatedness matrix (GRM) for heritability analysis, distinct ancestry subgroups had to be determined since the distribution of genetic relatedness is skewed when different ancestral subgroups are combined. Genetic ancestry of individuals is inferred from principal component analysis (PCA) of the genotyping data. **(A)** Partition around medoids (PAM) clustering yielded four optimal ancestry clusters consistent with the self-reported race and ethnicity of TCGA subjects: EUR, European ancestry (n=7,813), ASIAN, Asian ancestry (n=570), AFR, African ancestry (n=863), AMR American ancestry (n=209). **(B)** The first three principal components explaining the largest fraction of the variance in genotypes, PC1 vs. PC2 and PC vs. PC3 are shown with individuals color coded by ancestry group. **(C)** Analysis within each ancestry cluster identifies 33 unique immune traits with nominally significant level of heritability (LRT *p* < 0.05): 28 in the EUR cluster, two in the ASIAN cluster, and four in the AFR cluster. The AMR cluster is insufficiently powered. The corresponding heritability estimates with standard errors are shown for each of the 33 traits within each ancestry cluster with immune trait categories and corresponding immune trait modules annotated by the color bar.

**Supplementary Figure related to Figure 3.**
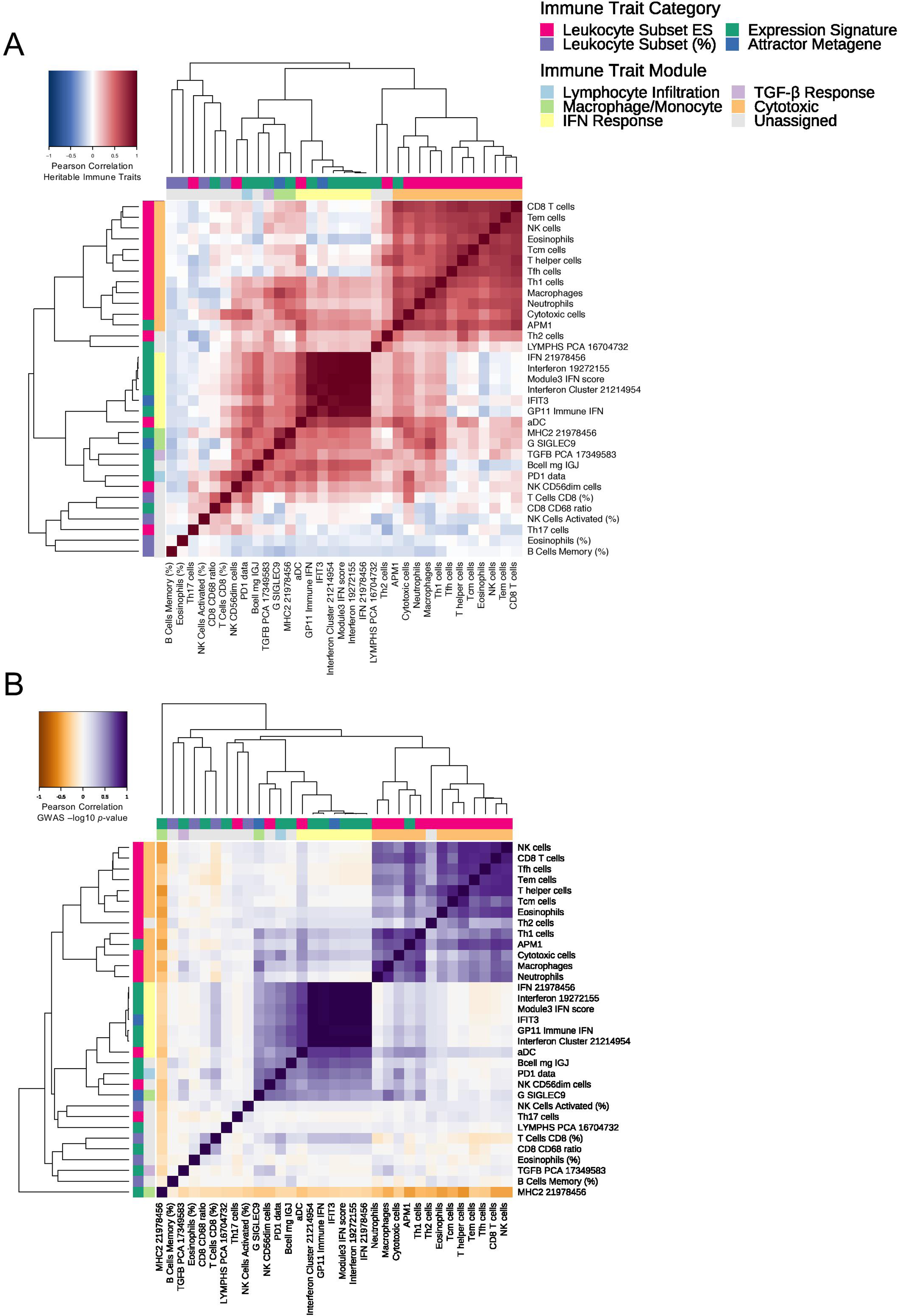
Characterization of GWAS results. Genome-wide association studies (GWAS) is performed in the 33 immune traits showing nominally significant genome-wide heritability (LRT *p* < 0.05) in at least one of the ancestry groups. **(A)** Heatmap of the pairwise Pearson correlation matrix of the phenotypic values. **(B)** Heatmap of the pairwise Pearson correlation matrix of the GWAS -log10 *p* of the immune traits. Immune trait modules and corresponding immune trait categories are annotated by the color bar.

**Supplementary Figure related to Figure 4.**
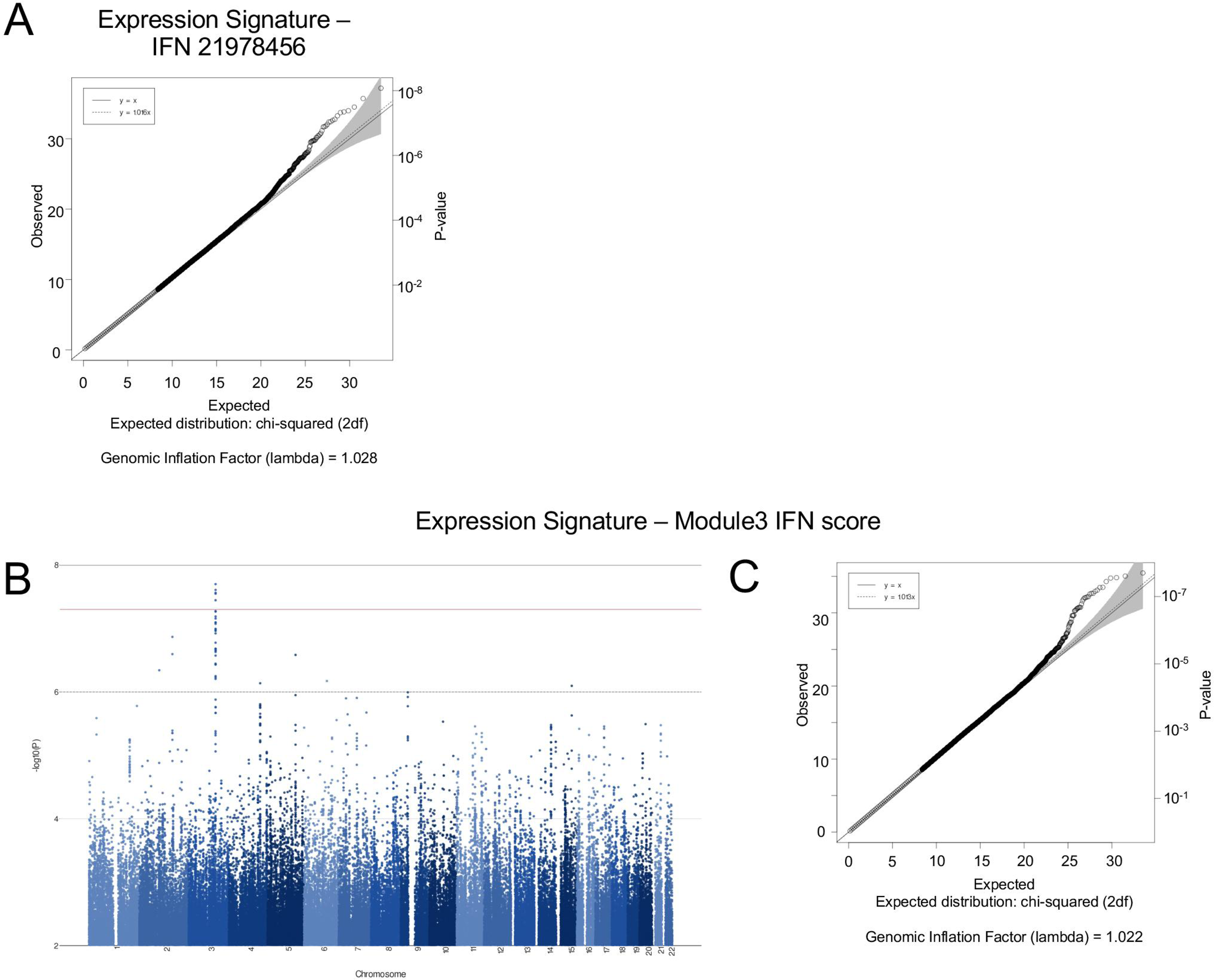
GWAS hits associated with Interferon-related signatures. Genome-wide association studies (GWAS) results for Interferon-related signatures. **(A)** QQ plot of IFN 21978456 GWAS results showing deviation of the observed *p* from the expected distribution from a theoretical χ^2^ distribution. Genomic inflation factor, lambda, is calculated. **(B)** Manhattan plot and **(C)** QQ plot from a second representative Interferon signature, Module3 IFN score which has the most significant TMEM108 associations (*p* < 5×10^−8^).

**Supplementary Figure related to Figure 5.**
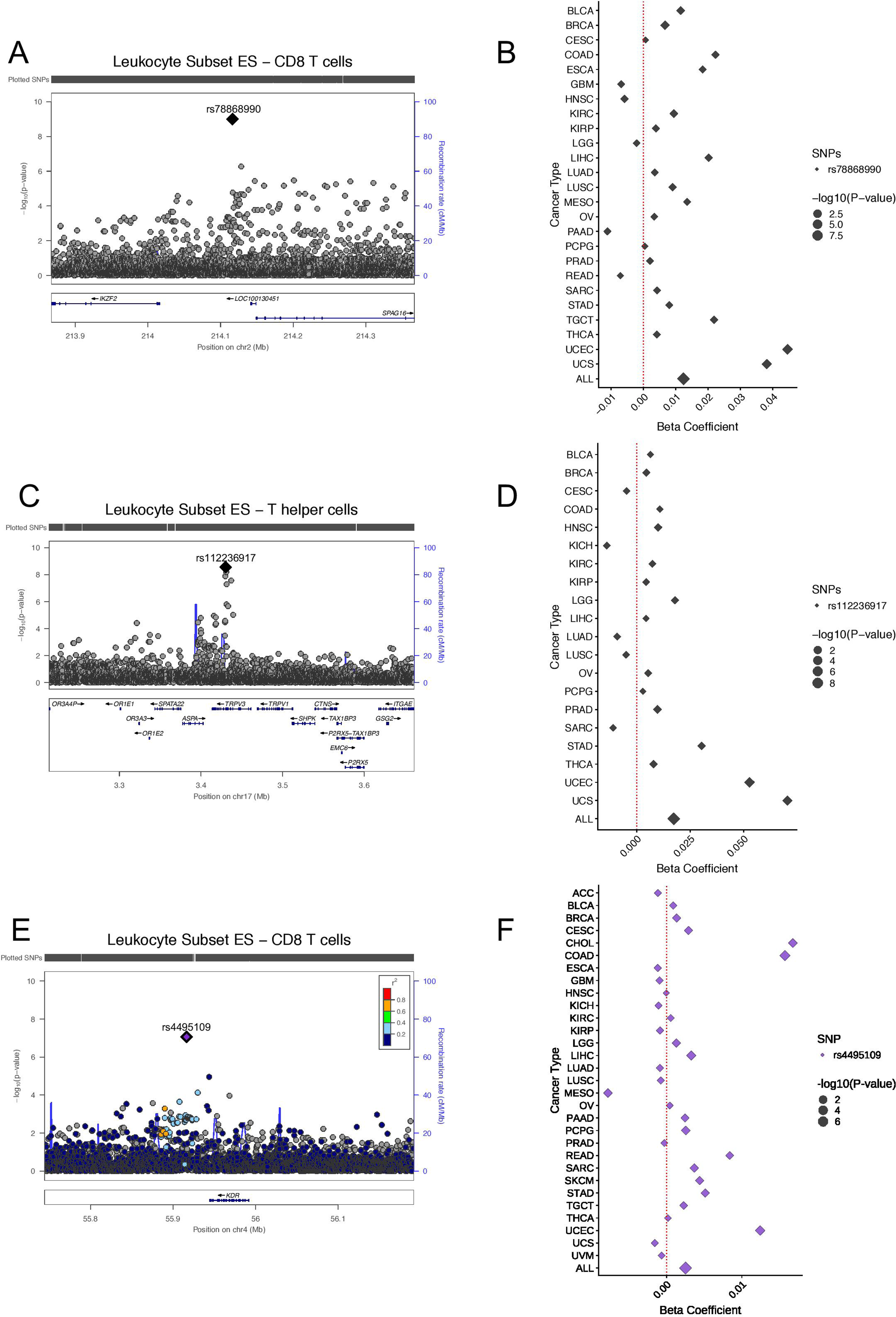
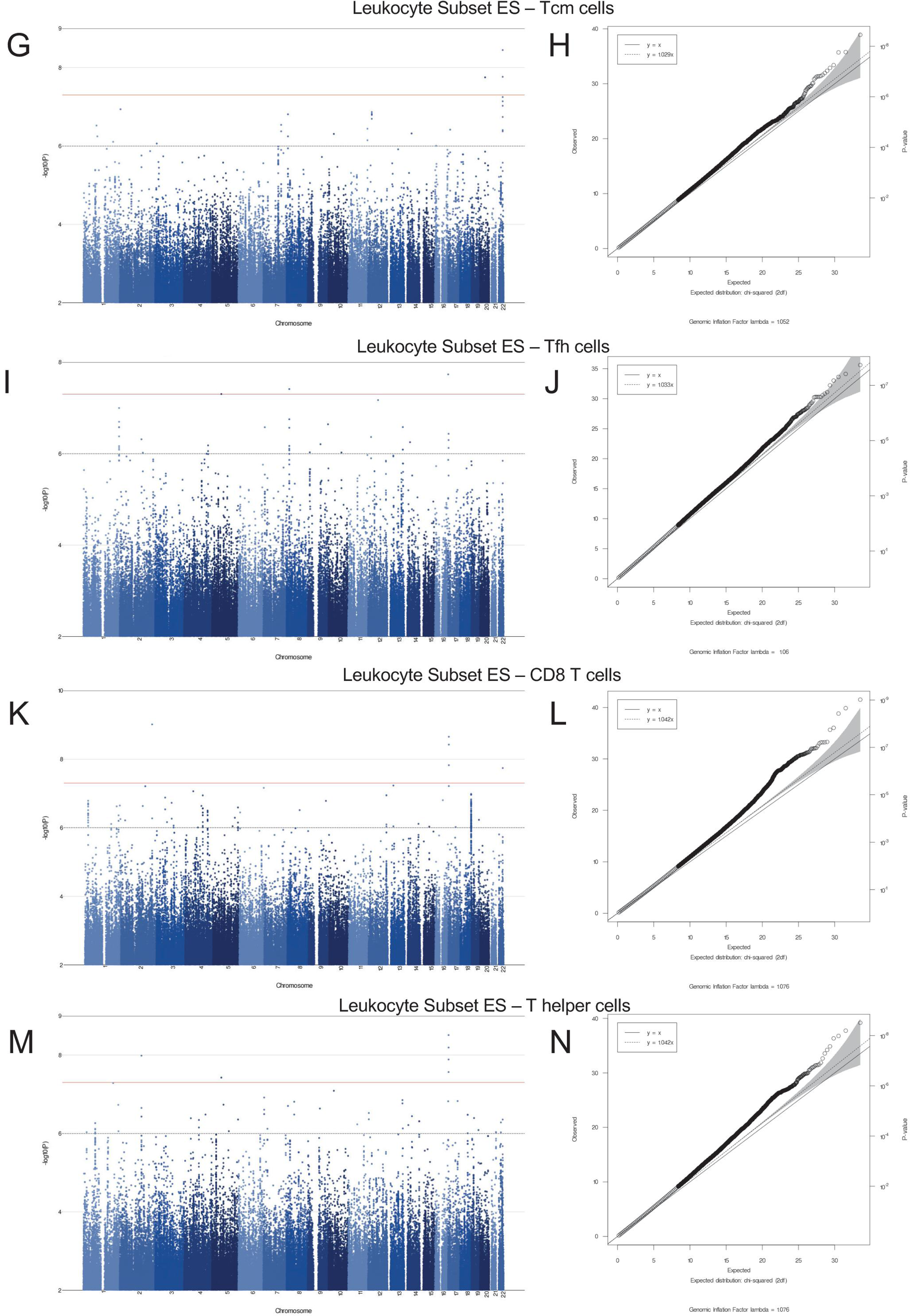
Germline variants associated with T-cell subset enrichment. Additional loci with multiple hits across multiple traits in the T-cell subset-dominant cluster. **(A)** Three associations involving two SNPs on chr 2 and CD8 T, and Cytotoxic cells mapping to in proximity to the SPAG16 locus are represented in a locus zoom plot. These SNPs are within 100KB of the *IKZF2* locus **(B)** Forest plot of Beta coefficients from association tests within each cancer type compared to the pan-cancer result for SNP rs78868990 in CD8 T cells. **(C)** Eleven associations involving 4 SNPs on chr 17 and T-helper, CD8 T cells, and T-follicular helper cells mapping to the TRPV3 locus are represented in a locus zoom plot. **(D)** Forest plot of Beta coefficients from association tests within each cancer type compared to the pan-cancer result for SNP rs112236917 in T-helper cells. **(E)** An association involving a SNPs on chr 4 and CD8 T cells mapping in proximity to the KDR locus are represented in a locus zoom plot. **(F)** Forest plot of Beta coefficients from association tests within each cancer type compared to the pan-cancer result for SNP rs4495109 in CD8 T cells. Within cancer association tests are run with age, gender and PC1-7 as covariates, except in CESC, OV, PRAD, TGCT, UCEC and UCS where only age and PC1-7 are used. Manhattan and QQ plots of GWAS results for immune traits in the T-cell subset-dominant cluster. (**G, I, K**, and **M**) Individual Manhattan plots and (**H, J, L**, and **N**) QQ plots of GWAS results for T-central memory, T-follicular helper, T-helper, and CD8 T cells.

**Supplementary Figure related to Figure 6.**
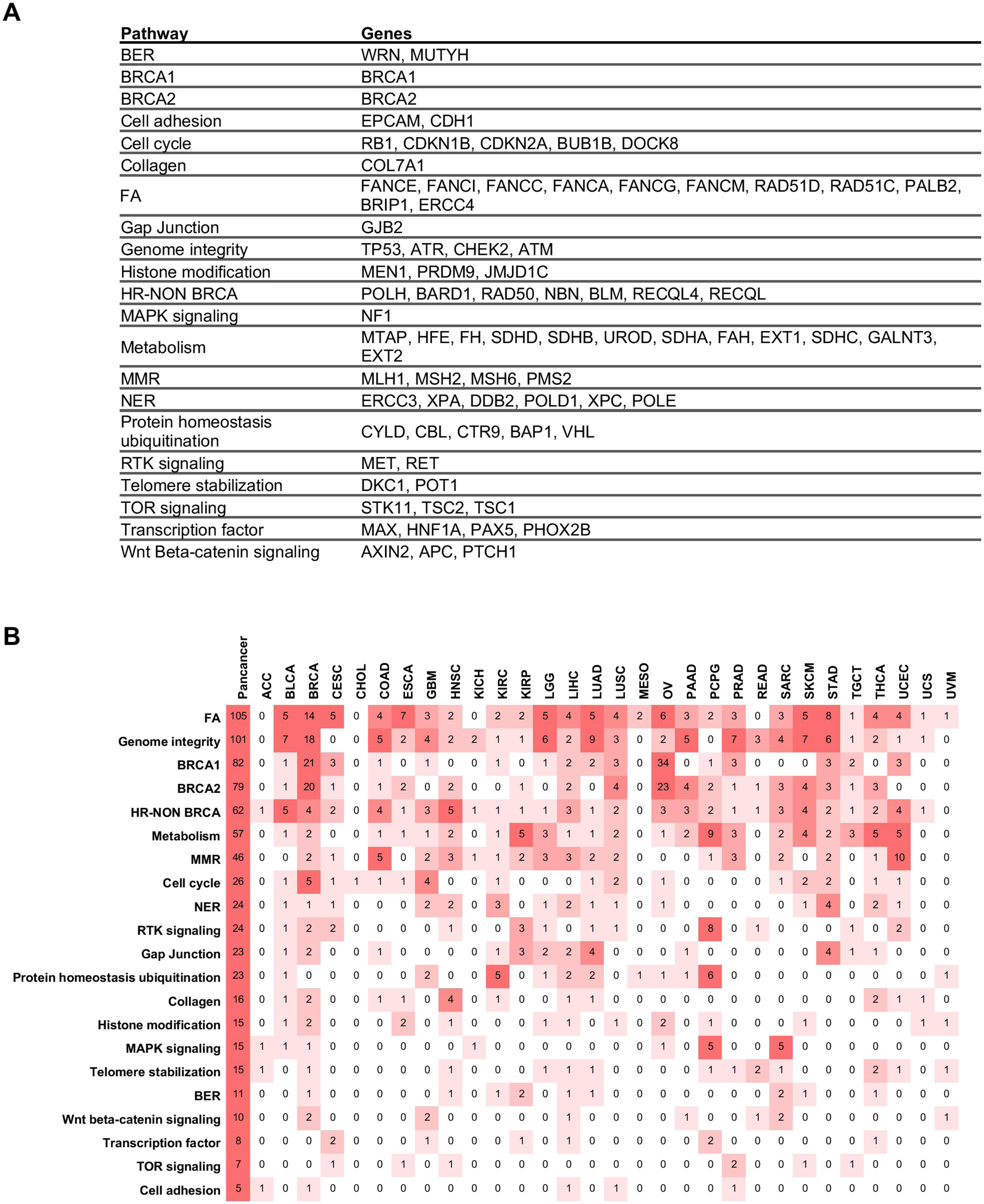
Overview of germline mutations by pathway and cancer type and associations between pathogenic and likely pathogenic germline variants and DNA-related somatic alterations. **(A)** Genes annotated by functional categories **(B)** Overview of total number of germline pathogenic or likely pathogenic mutations per cancer type grouped by functional categories used for the analyses. **(C)** Suggestive associations (*p* < 0.005 and FDR *p* ≤ 0.25) between rare genetic pathogenic or likely pathogenic variants extracted from whole-exome data (Huang KL, et al., *Cell*, 2018) grouped by curated mutually exclusive functional categories (*left nodes*) and DNA-related somatic alteration (*right nodes*) as identified in pan-cancer regression models adjusted for cancer type, age, sex, and PC1-7. Significant associations (*p* < 0.005 and FDR *p* < 0.1) are highlighted with blue dots. Within cancer, association tests are run with age, sex and PC1-7 as covariates, except in CESC, OV, PRAD, TGCT, UCEC and UCS where only age and PC1-7 are used. The adjusted-*p* for the non-silent mutation rate are also shown. Germline genotypic variables with a number of events lower than five across cancers were excluded from the analysis. Beta coefficients and significance level are visualized pan-cancer and per cancer (*right side*). Significance per cancer was only calculated when number of events was greater than two. The Beta coefficient is shown irrespectively of the significance and number of events. Beta: beta coefficient. Pan: *p* value pan-cancer. Pan+Nonsilent: *p* value pan-cancer adjusted for non-silent mutation rate. **(D)** Values of representative DNA-related traits (mean centered by cancer type) are displayed across samples with mutations in genes related to the defined functional categories. For indicated variables, log10(x+1) transformation was applied before values were mean centered by cancer type. BER: base excision repair; FA: Fanconi Anemia; HR NON BRCA: homologous recombination excluding *BRCA1* and *BRCA2*; MMR: mismatch repair; NER: nucleotide excision repair.

